# Multivariate analysis reveals a generalizable human electrophysiological signature of working memory load

**DOI:** 10.1101/2020.06.04.135053

**Authors:** Kirsten C.S. Adam, Edward K. Vogel, Edward Awh

**Affiliations:** Department of Psychology, University of California San Diego, La Jolla, CA 92093; Institute for Neural Computation, University of California San Diego, La Jolla, CA 92093; Grossman Institute for Neuroscience, Quantitative Biology, and Human Behavior, University of Chicago, Chicago, IL, 60637; Department of Psychology, University of Chicago, Chicago, IL, 60637; Institute for Mind and Biology, University of Chicago, Chicago, IL, 60637

**Keywords:** working memory, EEG, classification

## Abstract

Working memory (WM) is an online memory system that is critical for holding information in a rapidly accessible state during ongoing cognitive processing. Thus, there is strong value in methods that provide a temporally-resolved index of WM load. While univariate EEG signals have been identified that vary with WM load, recent advances in multivariate analytic approaches suggest that there may be rich sources of information that do not generate reliable univariate signatures. Here, using data from 4 published studies (*n* = 286 and >250,000 trials), we demonstrate that multivariate analysis of EEG voltage topography provides a sensitive index of the number of items stored in WM that generalizes to novel human observers. Moreover, multivariate load detection (“mvLoad”) can provide robust information at the single-trial level, exceeding the sensitivity of extant univariate approaches. We show that this method tracks WM load in a manner that is (1) independent of the spatial position of the memoranda, (2) precise enough to differentiate item-by-item increments in the number of stored items, (3) generalizable across distinct tasks and stimulus displays and (4) correlated with individual differences in WM behavior. Thus, this approach provides a powerful complement to univariate analytic approaches, enabling temporally-resolved tracking of online memory storage in humans.

## Introduction

Working memory serves as a critical interface between perception, memory, and action. Given the critical role of working memory in complex cognition, much prior work has been dedicated to identifying measures of the human electroencephalogram (EEG) signal that track working memory load in near real-time. The most widely-used of these measures is the contralateral delay activity (Vogel and Machizawa, 2004; Vogel et al., 2005), though other univariate measures such as suppressed alpha power and a sustained negative slow-wave have also been identified (Fukuda et al., 2015a, 2016). Tracking online working memory storage is critical for testing working memory’s role as an interface between varied cognitive demands. Studies taking advantage of univariate measures like the contralateral delay activity (CDA) have demonstrated filtering within working memory (Vogel et al., 2005), the role of working memory in guiding visual search (Emrich et al., 2009; Carlisle et al., 2011; Olivers et al., 2011; Woodman and Arita, 2011), the role of working memory in buffering retrieval from long-term memory (Fukuda and Woodman, 2017), and the role of existing long-term memories in shaping working memory encoding (Xie and Zhang, 2018). In cases where behavior is equivocal, neural measures are critical for disentangling competing explanations of an observed behavioral pattern. For example, unobtrusively monitoring the CDA during a typical visual search task revealed that search templates initially held working memory are moved to long-term memory with experience (Carlisle et al., 2011). Given the clear utility of near-real-time measures of working memory load, we present a novel analytic approach that provides a strong leap forward in the search for more sensitive and precise measures of storage in working memory.

Although univariate measures have been very productive for tackling many important questions about how and when working memory resources are deployed, they may miss some important aspects of the memory signal. For example, in the domain of spatial attention, it has long been known that lateralized, univariate changes to alpha power (e.g., contralateral alpha suppression) can be used to track attention to the left versus right hemifield, and that the topography of alpha power is modulated by finer-grained manipulations of spatial position (e.g., Rihs et al., 2007). In this context, multivariate analysis of alpha topography has been shown to provide a spatially- and temporally-resolved index of covert attention that substantially improves the utility of this signal for covert tracking of spatial attention (Foster et al., 2017). In addition to expanding the utility of known univariate measures, multivariate tools allow us to track information that was previously opaque to univariate analysis. For example, recent work has shown that motion direction of a dot cloud (Bae and Luck, 2019a) and a single remembered orientation (Wolff et al., 2015, 2017; Bae and Luck, 2018, 2019b) can be decoded from the topography of event-related potentials (ERPs; time-locked to stimulus or memory array onset), despite the absence of clear univariate signals that track this information.

Here, we show that a similar multivariate approach (“Multivariate load detection”, or “mvLoad”) enables tracking of online memory load in a sensitive and temporally-resolved fashion. Note, throughout the paper we define working memory load as the increasing amount of information held in mind with increasing memory set size. Although there is an ongoing debate about the format of mnemonic representations (e.g., item-based versus a flexible resource; Bays, 2018; Hakim et al., 2019), we do not directly address this issue here. In 3 experiments, we demonstrate that we can predict working memory set size from ERPs of small groups of trials (time-locked to memory array onset) and even with single trials of EEG data. Further analyses demonstrate that this multivariate decoding signal has the expected profile of a working memory signal (e.g., modulated by working memory task demands; shows higher confusability for supra-capacity working memory loads) and carries promise for future cross-subject, cross-experiment decoding applications.

## Materials and Methods

### Overview of Datasets

We used four previously published datasets to examine whether and why we can decode working memory load from the electroencephalogram (EEG) signal. Further methodological details about the participants, tasks, and data acquisition can be found in each of the original published papers (Unsworth et al., 2014, 2015; Fukuda et al., 2015a, 2015b; Hakim et al., 2019). Some high-level information about the studies is provided in Table 1 and below, and additional information about pre-processing of the EEG data is available in the Supplemental Methods. Figure 1 shows examples of the stimuli and task procedures. Data for all experiments are freely available the Open Science Framework at https://osf.io/6jkqu/

**Table 1.** Overview of datasets

| Exp. | Paper | Task | Set Sizes | Total trials | Subjects (n) |
| --- | --- | --- | --- | --- | --- |
| 1 | Unsworth et al., 2014; 2015 | Lateralized change detection | 2, 6 | 116,101 | 152 |
| 2a | Fukuda, Mance, Vogel 2015 | Lateralized & Whole-field change detection | 1-4, 6, 8 | 23,951 | 30 |
| 2b | Fukuda, Woodman, Vogel 2015 | Lateralized change detection | 1-8 | 49,916 | 31 |
| 3 | Hakim et al. 2019 | Lateralized change detection; Lateralized attention task | 2, 4 | 102,358 | 73* |

\*73 unique subjects participated in one or more of the 4 sub-experiments (total of 97 sessions).

**Figure 1.**
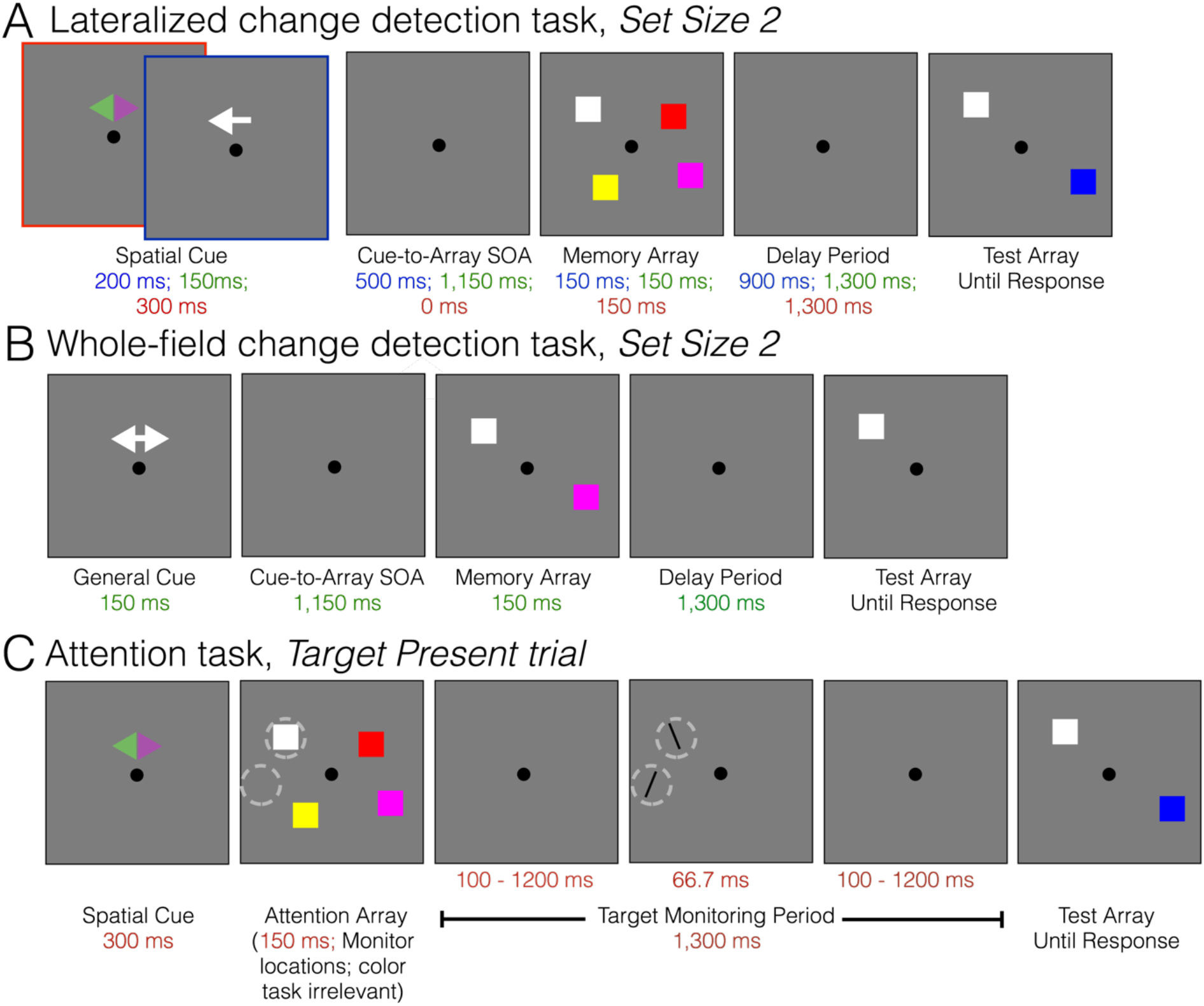
Task schematics for a typical Set Size 2 trial. (A) Lateralized changed detection task used in Exps. 1, 2a, 2b, and 3. Participants are first cued to one side of space (Red cue box shows symbolic cue used in Exp 3, e.g., attend green side; Blue cue box shows spatial cue used in other Exps.). Participants remember the items on the cued side, and are tested on one of the items. Blue text shows task timing for Exps. 1 and 2b; Green text shows timing for Exp. 2a; Red text shows task timing for Exp 3. (B) Whole-field change detection task used in Exp 2a. Participants are given a non-spatial cue that the trial is upcoming, remember all items on the screen across a delay, and are tested on one of the items. (C) Attention task used in Exp 3. This task serves as a control for working memory task demands. Participants are cued to one side of the display. Rather than remember the colors of the squares, participants are asked to pay attention to the spatial positions occupied by the squares. They monitor these positions across a target monitoring period. A target (tilted line at the same location as one of the item positions) and/or distractor (tilted line at a foil location where no item appeared) appear on 25% of trials. For illustrative purposes only, the locations of the target and distractor lines are circled. The colored squares shown in the test array are task-irrelevant (visual control for WM condition); the array indicates when participants should make their response about the target.

#### Experiment 1

Participants performed two conditions of a lateralized change detection task (Unsworth et al., 2014, 2015); EEG data were collected from 20 passive electrodes. In one condition, participants remembered the colors of items (color change detection; set size 2 and 6). In the other condition, participants remembered the shapes of items (shape change detection, set size 2 and 6). Here, we collapse across the color and shape conditions (i.e., just examining set size 2 versus 6 overall). See Figure S1 for confirmation that the general results hold for separately analyzed color and shape conditions. The full dataset includes 183 participants, with 200 trials per sub-condition (e.g., “shape set size 2”). Participants were analyzed if they had at minimum 160 trials per sub-condition after artifact rejection, leaving 152 subjects for analysis (M = 382 trials per set size after collapsing across color and shape).

#### Experiment 2a

Participants completed two conditions of a change detection task: lateralized change detection and whole-field change detection (Fukuda et al., 2015a); EEG data were collected from 20 passive electrodes. Participants (*n* = 29) completed 80 trials per set size (1-4, 6, and 8) in each condition (M = 68.8 trials per set size in each task condition).

#### Experiment 2b

Participants (*n* = 31) completed a lateralized change detection task (Fukuda et al., 2015b); EEG data were collected from 20 passive electrodes. There were eight set size conditions (1-8; M = 201.3 trials per set size).

#### Experiment 3

The full dataset includes 97 sessions from 4 sub-experiments (some additional sessions were collected but excluded according to the artifact rejection criteria detailed in (Unsworth et al., 2014, 2015; Fukuda et al., 2015a, 2015b; Hakim et al., 2019)EEG data were collected from 32 active electrodes. In published work, we demonstrated no major differences between these 4 sub-experiments and we performed analyses collapsed across all 4 sub-experiments (Hakim et al., 2019). We likewise combined data across all 4 sub-experiments here. All participants completed two key task conditions: (1) a lateralized working memory task, and (2) a lateralized attention task. In some sub-experiments, the lateralized working memory task employed color memoranda and in other cases the lateralized working memory task employed spatial memoranda. In all cases, there were two set sizes (2 or 4 items) in each task (M = 263.8 trials per set size in each condition). Although participants could complete each sub-experiment only once, some participants completed multiple sub-experiments. This resulted in a total of 73 unique subjects. If a participant completed more than one sub-experiment, we averaged their behavioral and classification results across sessions so that each participant was equally represented in the full dataset.

### Tasks

#### Lateralized change detection

On each trial, participants are first cued to attend one hemifield (left or right) with a brief spatial cue. After the cue, there is a blank interval (“cue-to-array SOA”), and then the memory array appears. The memory array consists of brightly colored squares drawn in both hemifields. Participants are instructed to remember only the colored squares in the cued hemifield across a blank delay period. At test, a single colored square is presented at one of the remembered locations. On 50% of trials (“same trials”), the test square is the same color as the item presented at that position. On the other 50% of trials (“change trials”), the test square is a different color from before. Participants press one of 2 keys to indicate whether the test square is the same color or has changed colors. The exact task timing varied slightly across datasets. Experiments 1 and 2b used a cue duration of 200 ms, cue-to-array SOA of 500 ms, memory array duration of 150 ms, and a delay period of 900 ms. Experiment 2a used a cue duration of 150 ms, cue-to-array SOA of 1,150 ms, memory array duration of 150 ms, and a delay period of 1150 ms. The working memory condition from Experiment 3 used a cue duration of 300 ms, cue-to-array SOA of 0 ms, array presentation of 150 ms, delay period of 1,300 ms, and a blank inter-trial interval of 750 ms. Experiment 3 is a combination of 4 sub-experiments reported in Hakim et al. 2019. The task events and timing are consistent across all 4 sub-experiments. In 2 sub-experiments participants remembered color (as described); in the other 2 sub-experiments, participants remembered the spatial position of items, and were tested with an item that was either at the same location (“same trial”) or with an item that appeared at a foil location a minimum of 1.5 objects’-width away from any of the remembered locations (“different trial”). Prior work revealed that these stimulus-specific differences did not greatly alter the contralateral delay activity, and that it was justified to collapse across these sub-experiments for further analysis(Hakim et al., 2019).

#### Whole-field change detection

Experiment 2a used a whole-field version of the change detection task. This task is very similar to the lateralized change detection task, except there is no spatial cue. Instead, participants receive a task-general cue (e.g., a double-sided arrow that does not indicate a side to attend but gives a temporal warning that the memory array is coming). Participants remember all items from the array, and to-be-remembered items are presented in both the left and right hemifields (i.e., “whole-field”). As before, participants remember colors across a delay and are probed on one item, and report whether the probed item is the same as or different from the remembered item. Exp 2a used a cue duration of 150 ms, cue-to-array SOA of 1150 ms, memory array duration of 150 ms, and a delay period of 1150 ms.

#### Lateralized attention task

In Experiment 3, the relative need for working memory task demands was manipulated. Participants viewed identical stimuli as in the lateralized change detection task described above, but they were given different task instructions which could be achieved with sustained spatial attention. This task used identical stimuli and task timing as a lateralized change detection task, but asked participants to perform a sustained attention task, rather than a working memory task. This allows us to compare mental processes for “attention” and “working memory” tasks while holding visual stimulation and task timing constant. In prior work (Hakim et al., 2019), we showed that the contralateral delay activity was present in the working lateralized change detection task but not in the lateralized attention task, indicating that the CDA is associated with working memory task demands. In the task, participants are first cued to one visual hemifield (e.g., attend the hemifield indicated by the green side of a double-sided arrow; colors counterbalanced across participants). Rather than remembering the colors and/or locations of the items in the “memory array” (e.g., array of colored squares), participants were instead instructed to maintain their spatial attention to the positions occupied by the items. Participants maintained spatial attention to the positions during a blank array in order to detect and discriminate a rare target (titled line) that briefly appeared (66.67 ms) at one of the attended positions on 25% of trials. Note, there was always one target (line that occupied one of the same spatial positions that was cued). In addition, in 3 of 4 sub-experiments a distractor line was shown at a “foil” location.

During the attention task, the color and position of the item in the “test array” were task-irrelevant. Instead, the test array simply indicated the time when participants should make their response. The participants made one of three button presses: (1) target absent, (2) target present, top tilted left, (3) target present, top tilted right. Note, we discarded the 25% of target-present trials for analysis to avoid any potential physical display confounds. We analyzed only the 75% of trials where there was a fully blank delay period. As expected, in the working memory task participants stored slightly more items for set size 4 (M = 1.73) versus set size 2 trials M = 1.52; *p* = .003). Likewise, participants had poorer performance in the attention task when they monitored 4 locations (M = 78% correct) versus 2 locations (M = 82% correct; *p* = .01), see Hakim et al. (2019) for further discussion.

### EEG data acquisition

#### Experiments 1-2

Experiments 1, 2a, and 2b were collected from 20 passive tin electrodes (SA Instrumentation Co., San Diego, CA) mounted in an elastic cap (ElectroCap International, Eaton, OH). Electrode positions included International 10/20 sites F3, Fz, F4, T3, C3, Cz, C4, T4, P3, Pz, P4, T5, T6, O1, and O2 and five non-standard sites: OL midway between T5 and O1, OR midway between T6 and O2, PO3 midway between P3 and OL, PO4 midway between P4 and OR, POz midway between PO3 and PO4. Data were recorded with a left-mastoid reference and re-referenced offline to the algebraic average of the left and right mastoid. Horizontal electrooculogram (EOG) and vertical EOG were collected from 3 additional passive electrodes affixed to the face with stickers. Trials containing ocular artifacts, movement artifacts, or blocking were excluded from analyses.

#### Experiment 3

Experiment 3 was collected from 30 active Ag/AgCl electrodes (actiCHamp, Brain Products, Munich Germany) mounted in an elastic cap positioned according to the international 10-20 system (Fp1, Fp2, F7, F8, F3, F4, Fz, FC5, FC6, FC1, FC2, C3, C4, Cz, CP5, CP6, CP1, CP2, P7, P8, P3, P4, Pz, PO7, PO8, PO3, PO4, O1, O2, Oz). Two additional active electrodes were affixed with stickers to the left and right mastoids, and a ground electrode was placed at position Fpz. Data were referenced online to the right mastoid and re-referenced offline to the algebraic average of the left and right mastoids. Passive electrodes (HEOG, VEOG) and eye tracking were used to monitor eye movements and blinks. Trials containing ocular artifacts, movement artifacts, or blocking were excluded from analyses.

### Classification and significance testing

#### Single-trial classification (within-subject)

Classification was performed within a subject, on single trials, and within a given time window on raw, baselined EEG data (Exp 1). We divided each trial into 50-ms windows and calculated the average voltage for each electrode within this window (e.g., 20 electrodes = 20 predictors). Classification was performed separately within each time point using a linear discriminant classifier (“classify.m”, with option ‘diagLinear’ to use the diagonal covariance matrix estimate; MATLAB, MathWorks, Natick MA). We chose to use a relatively simple linear classifier to first demonstrate our effect, but our results are expected to also generalize to other classification methods such as support vector machines (SVM; Figure S2). We performed 100 iterations of the classification analysis at each point; on each iteration, we randomly assigned 2/3 of the trials to an independent training set and 1/3 of the trials to a held-out test set. We also confirmed that the trial voltage distributions on each iteration of the analysis were approximately normally distributed (Figure S3). A schematic of the classification procedure is shown in Figure 2A. In all classification procedures, the number of trials per set size was balanced in both the training and test sets.

**Figure 2.**
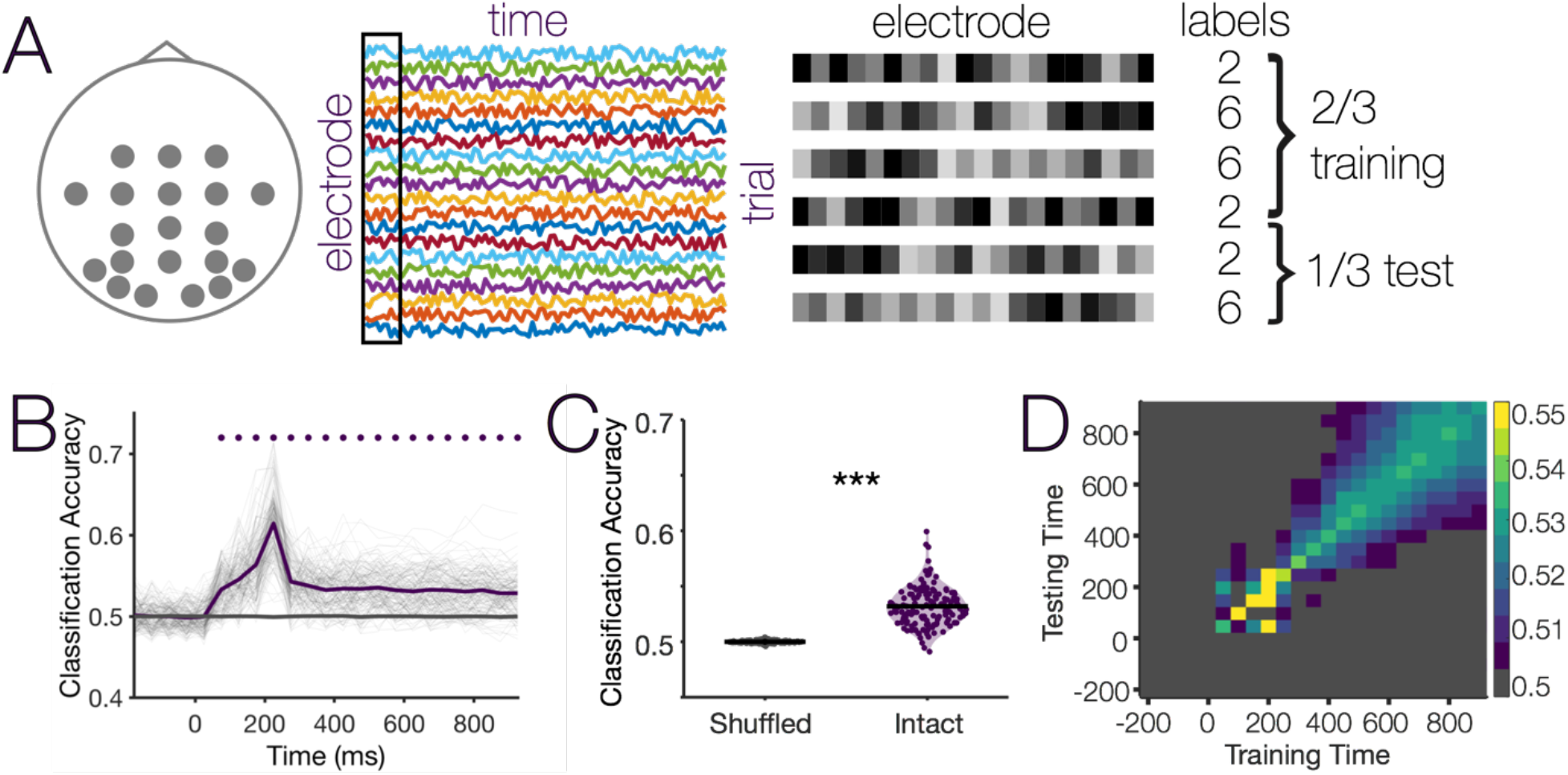
Single-trial decoding of working memory set size in Experiment 1. (A) Schematic of single-trial decoding approach. Decoding was performed within a participant, separately at each time bin (1 average value per electrode for a 50 ms time bin) using data from all electrodes (Data: *t* trials x 20 electrodes; Labels: 1 x *t* trials). On each iteration of the analysis, we picked a random 2/3 of trials to serve as a training dataset, and the remaining 1/3 of trials were used as a testing dataset. (B) Single trial decoding performance over time. Expected chance is 50%; dots indicate Bonferroni-corrected *p* < .001. (C) Average single trial decoding performance during the delay period (400 – 1000 ms; *** *p* < .001). (D) Cross-temporal generalization of classification performance (training and testing across different time bins in the trial). Gray indicates that the pixel did not survive the cluster-based permutation test.

#### Mini-block classification (within-subject)

Instead of performing classification on single trials, we averaged together groups of like trials (i.e., the same set size condition) into “mini-blocks”, and we shuffled the assignment of trials to different mini-blocks across 100 iterations (Exps. 2 & 3). Classification was still performed using the same linear classification routine, training on 2/3 of mini-blocks and testing on a held-out 1/3 of mini-blocks on each iteration. Also note, this classification function handles both binary and multi-class classification, so the same general classification methods were used for both Experiments 1 and 3 (binary) as well as Experiment 2 (multi-class). To assess whether classification generalized across tasks (Exps 2 & 3), we used the same method except we trained on 2/3 of mini-blocks from one task (e.g., lateralized change detection) and tested on 1/3 of mini-blocks from the other task (e.g., wholefield change detection).

#### Mini-block classification (across-subject)

To test the generalizability of the classification signal across subjects, we performed a leave-1-subject-out analysis (Exp 2). To do so, we blocked trials in the same way as in the within-subjects version of the mini-block analysis. We trained the classifier on all data on a random 2/3 of mini-blocks from *n*-1 subjects and tested data on a random 1/3 of mini-blocks from 1 held-out subject. For each held-out subject, we ran 100 iterations of randomly assigning individual trials to mini-blocks in both the training and test sets.

#### Statistical tests

In the standard within-subject classification analyses, we trained and tested on data from the same time bin (e.g., train and test on data averaged from 0 – 50 ms). Significance of the time-course of overall classification was assessed via Bonferroni-corrected *t*-tests (one-sided *t-*tests when comparing to chance level, as we would not expect to find meaningfully below-chance values; two-sided tests when comparing between conditions). To assess the generalizability of the signal across time points, we also performed a cross-temporal analysis where we trained and tested on all possible combinations of time-points (e.g., train on the first time point, test on all other time points). Significance of the generalizability of decoding was assessed via a cluster-based permutation test statistic, based on comparing significant clusters of adjacent significant *t-*values to a permuted distribution (Maris and Oostenveld, 2007), using a subject-wise permutation function (1,000 iterations) adapted from Fahrenfort et al. (2018). Significance was always assessed in comparison to empirical chance values (rather than to theoretical chance, see Combrisson & Jerbi, 2015); empirical chance was estimated by repeating the same classification analysis using randomly shuffled training labels.

## Results

### Experiment 1

#### Single-trial classification predicts working memory load in a large sample

Using a large sample (n = 152, *trials* = 116,101) and a linear classifier on raw EEG amplitudes from single trials (Figure 2A), we could predict working memory load (set size 2 versus set size 6) in a sustained fashion throughout the delay period (Figure 2B; dots indicate *p* < 1×10^−5^, Bonferroni-corrected for 23 time-bins). We could classify set size quite early in the trial (50-100 ms time bin), although this early classification could be due to a physical display difference between the two and six item arrays. Importantly, classification was sustained throughout the delay in the absence of any physical display differences. Mean decoding accuracy during the delay period (400 – 950 ms) was 53.2% (SD = 1.7%; Figure 2C), significantly above the chance level produced by giving the same classifier shuffled labels, *t*(151) = 22.5, *p* < 1 × 10^−49^, 95% CI [2.92%, 3.48%], Cohen’s *d* = 1.85. Finally, decoding generalized to other time-points beginning at the 400-450 ms time bin and lasting throughout the delay (Figure 2D; gray boxes indicate that the pixel did not survive a cluster-based permutation test). Notably, the decoding signal observed during encoding (100 - 300 ms) did not generalize throughout the delay period, indicating that it is unlikely that a sensory imbalance signal early in the trial drove the sustained delay period decoding.

#### Global versus lateralized contributions to decoding

Although our analysis shows that topography of voltage across electrodes predicted working memory load on a single trial basis, this simple decoding approach was blind to *lateralized* EEG signals that track working memory storage, such as contralateral delay activity (CDA) a well-documented electrophysiological marker of storage in visual working memory. Thus, the robust performance of our decoder could not be explained by contributions from CDA activity. Nevertheless, this leaves open the interesting question of whether load detection could be further improved by taking lateralized storage signals into account. To test this, we added 8 new predictors corresponding to single-trial paired difference waves (contralateral minus ipsilateral, e.g., PO8 minus PO7 for a “remember left” trial, PO7 minus PO8 for a “remember right” trial). Surprisingly, we found that adding lateralized predictors did not predict substantial additional variance beyond the single electrodes (Figure 3A). On its own, activity from just the 8 lateralized predictors likewise tracks working memory load in a sustained fashion throughout the delay period (*M* = 52.9%, *SD* = 1.7%, *p* < 1×10^−16^, *d* = 1.67), in line with many past demonstrations in the CDA literature. Including both single electrodes and the 8 lateralized predictors somewhat improved classification performance beyond the lateralized predictors alone (53.1% combined versus 52.9% lateralized alone, *p* < .001). Critically, however, a classifier combining both the 8 lateralized predictors and the 20 single electrode predictors did slightly *worse* than the 20 single electrode predictors alone (53.1% combined versus 53.2% single electrodes alone, *p* < .001). Further control analyses revealed that training and testing within a cued side (e.g., train and test on “remember right” trials) offered only marginal benefits over training and testing across cued sides (e.g., train on “remember right” test on “remember left”), see Figure S4.

**Figure 3.**
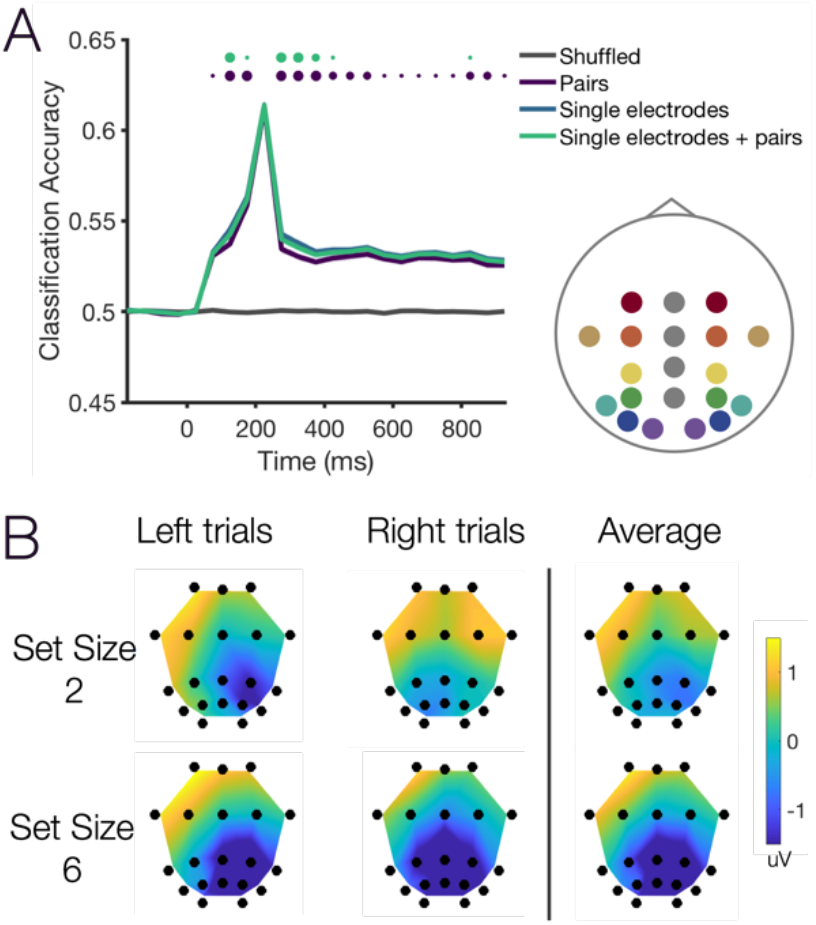
Adding lateralized predictors does not improve classification performance. (A) We created 8 additional predictors by taking the difference (contra – ipsi) of all matched pairs of lateralized pairs (“pairs”, color-coded in the inset diagram). Lateralized predictors did a good job of predicting working memory load, but did not improve classification above the level of the 20 single electrodes. In this figure, dots represent Bonferroni-corrected significance (small *p* <.05, medium *p* < .01, large *p* < 001). Purple dots represent the difference between single electrodes and pairs alone. Green dots represent the comparison between “single electrodes + pairs” and pairs alone. (B) Topographical plots of delay period activity for each set size condition, separated by side or averaged across both sides. Color scale is in microvolts (μV).

Although it is somewhat surprising that adding lateralized predictors did not improve decoding, topographic plots for each set size condition suggest that this failure to explain additional variance may be due to the coarse spatial distribution of the signal (Figure 3B) and/or to noisiness of using difference waves as predictors on a single trial basis. As adding the lateralized predictors failed to improve classification performance, we will continue to use the single electrode classifier (i.e., giving the classifier the raw voltage value at each electrode). Arguably, insensitivity to stimulus laterality makes the analysis more flexible and powerful, providing the potential to train and test classifiers across lateralized and non-lateralized working memory tasks; we provide an example of such cross-training in Exp 2A/B. In supplemental analyses, we examined decoding accuracy separately for individual electrodes and groups of electrodes (e.g., occipital alone, Figure S5), and we confirmed that decoding was not driven by a global signal alone (Figure S6). Together, the topography of voltage changes and the supplementary analyses suggest that broadly-distributed changes to voltage values (i.e., changes to the degree/extent of frontal positivity and changes to the degree/extent of posterior negativity) are likely driving overall decoding performance.

#### Improving decoding with mini-block classification

Although the prior analyses demonstrated that single-trial classification is extremely robust during the delay period (*d* = 1.85), single trials are noisy and thus classification accuracy is numerically low (∼53% given a chance level of 50%). To accommodate the lower numbers of trials and participants in some experiments, we tested whether averaging across small sub-sets of trials (“mini blocks”) would improve overall classification accuracy (Figure 4). Here, we performed the same basic analysis (training on 2/3 of data and testing on 1/3 of data, for 100 iterations of randomly assigning trials to training or testing). However, rather than training on single-trials, we averaged small groups of trials together to reduce noise of each instance given to the classifier. This “mini-block” procedure was quite effective at improving overall classification accuracy, both early and late in the trial (Figure 2A). During the encoding period (100-300 ms), classification accuracy improved monotonically with the number of trials per mini-block, *F*(1.1,155.7)^*^ = 2615, *p* < 1×10^−106^, η^2^_p_ = .95. In the peak sensory time bin (200- 250 ms), decoding accuracy reached as high as 89.4% (SD = 7.7%; 25 trial mini-blocks). During the delay period, classification accuracy likewise improved monotonically with the number of trials per mini-block, *F*(1.03,156.14) = 571.6, *p* = < 1×10^−54^, η^2^_p_ = .79, topping out at 63.9% (SD = 7.1%) with 25 trial mini-blocks (Figure 2B). Note, however, the number of trials that may be used in mini-blocks is limited by the number of available trials per condition. For consistency of comparisons across experiments, we will use 10-trial mini-blocks for further analyses. This will allow us to improve classification accuracy while still accommodating the varied numbers of trials per experiment (∼80 – 400 trials per set size).

**Figure 4.**
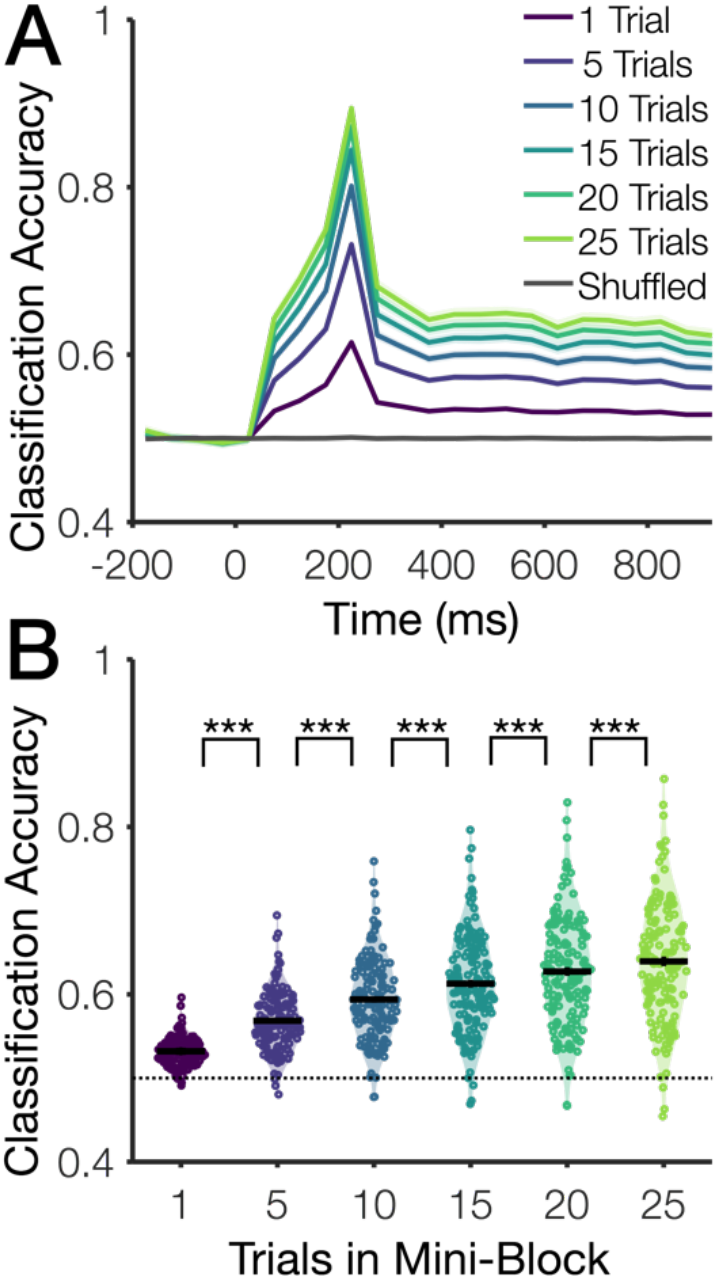
Decoding accuracy by number of trials in classification “mini-blocks”. We examined changes to decoding accuracy for training and testing on single trials versus training and testing on averaged values from small groups of trials (e.g., a “mini block” of five Set Size 2 trials). (A) Classifier accuracy over time for single trial decoding and mini-block decoding of various block sizes. Shaded error bars indicate ±1 SEM. (B) Mean classification accuracy during the delay period (400 ms – end of delay). *** *p* < .001 (Bonferroni corrected, 5 comparisons).

### Experiment 2

#### Decoding differentiates item-by-item increments in load

In Exp 1, we demonstrated that we can discriminate between two memory load conditions (set size 2 vs. set size 6) using the multivariate EEG signal across electrodes (training and testing the classifier within a subject). In Exp 2A and 2B, we tested whether this within-subject classification signal is sensitive to finer-grained set size manipulations. In Exp 2A, participants performed two working memory tasks (lateralized and whole-field) with 6 set sizes (1-4, 6, and 8); In Exp 2B, participants performed a lateralized working memory task with 8 set sizes (1-8). As shown in Figure 5A, we could robustly predict set size in a sustained fashion throughout the delay period (all *p’*s < 1×10^−8^, Cohen’s *d =* 1.55, 1.49, and 2.27 for panels left to right in Figure 5A). As in Exp 1, this classification signal sustained through the end of the delay and generalized to all later time bins beginning mid-way through the delay (650-700 ms, 600-650 ms, and 600-650 ms, for the panels shown left to right in Figure 5B).

**Figure 5.**
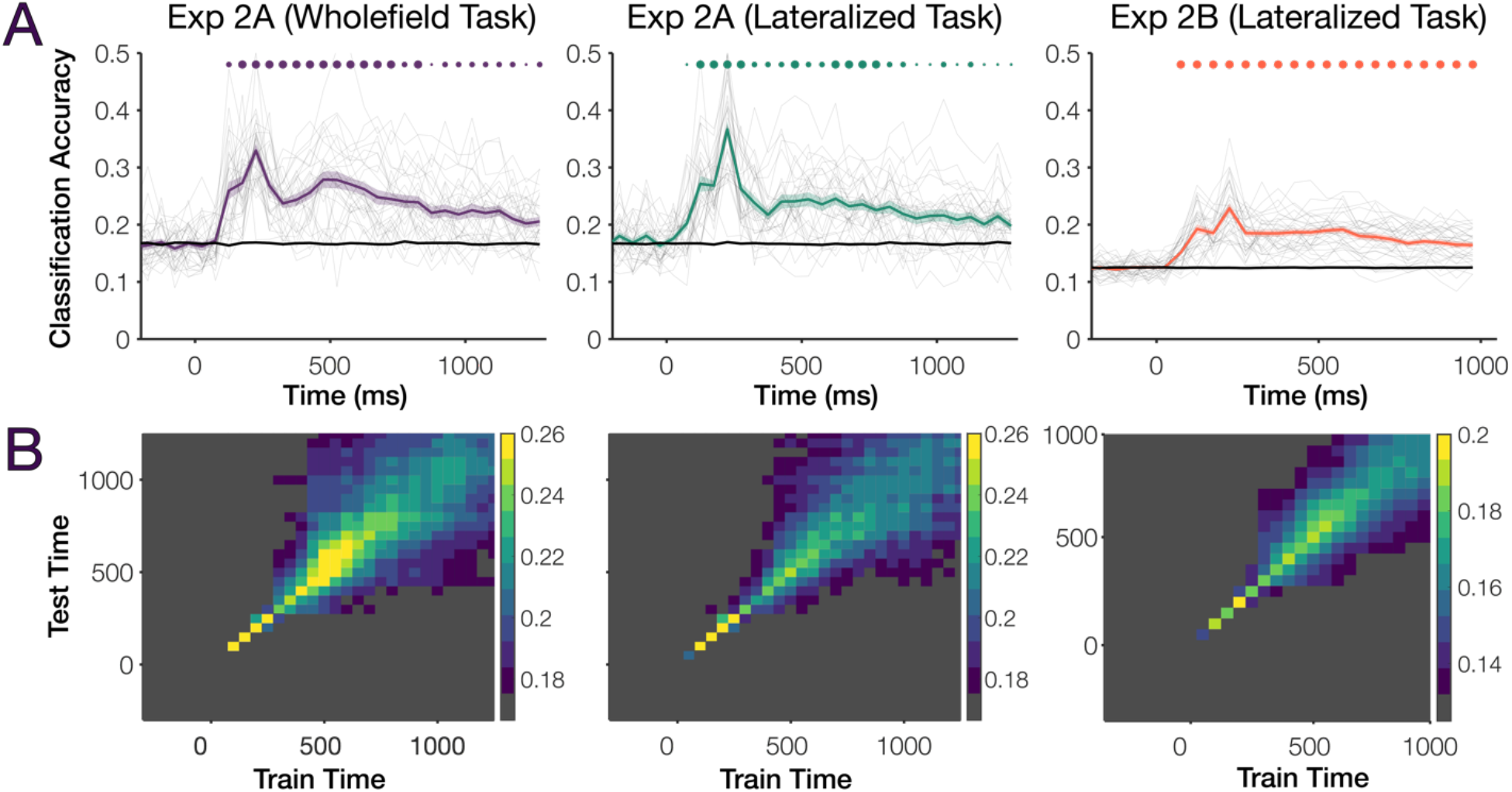
Decoding accuracy in Experiments 2A and 2B. (A) Classification accuracy over time for Exps. 2A and 2B. Chance level is 1/6 in Exp 2A and 1/8 in Exp 2B (black line depicts empirical chance from shuffled analysis). Shaded error bars represent ±1 SEM. Transparent gray lines represent individual subjects. Dots indicate Bonferroni-corrected significance of each 50 ms time-bin (small dots, *p* < .05, medium, *p* < .01, large, *p* < .001). (B) Cross-temporal generalization of classification performance (training and testing across different time bins in the trial) for Exps. 2A and 2B. Gray indicates that the pixel did not survive the cluster-based permutation test.

In addition to examining overall decoding, the inclusion of more set sizes in Experiments 2A and 2B provided the chance to look at classifier errors and discriminability of set size. Figure 6A and 6B shows confusion matrices for the encoding period (100-300 ms) and delay period (400 ms to end of delay) in Exps. 2A and 2B. Much prior work on univariate neural signatures of working memory maintenance (e.g., (Todd and Marois, 2004; Vogel and Machizawa, 2004; Xu and Chun, 2006) has found that, consistent with a capacity limit of 3-4 items, univariate measures increase from set sizes 1 to 3, but reach an asymptote around 3-4 items (but see Bays, 2018). Given this key signature of univariate working memory measures, we predicted that delay period decoding, but not sensory period decoding, should show particularly poor or no discriminability amongst larger set sizes. To test this, we compared confusion matrix discriminability amongst the lower set sizes (1-3 in Exp 2A, 1-4 in Exp 2B) versus discriminability amongst higher set sizes (4,6,8 in Exp 2A, 5-8 in Exp 2B) in both the encoding period and delay period. Discriminability was quantified as the difference between the true category value (e.g., proportion of the times the classifier chose set size 1 when the true value was 1) and the mean of the incorrect values (e.g., how often the classifier instead chose other low set sizes 2 or 3). As we predicted for the delay period signal, we observed significantly higher decoding amongst low set sizes than high set sizes for all 3 experiments (*p* < .001) and we observed a null effect for discriminability amongst high set sizes during the delay in all 3 experiments (*p* > .10). We likewise observed higher discriminability amongst low set sizes than high set sizes during the sensory period (100-300 ms, all p’s < .01) but significant encoding period decoding for both high and low set sizes (all p’s < .001). For uncluttered visualization, here we have shown confusion matrices as color-scales; Please see Figure S7 in the supplemental for numerical values in each cell of the confusion matrix. These effects was similar when we instead used many pairwise classifiers to test which set sizes were discriminable from one another during the delay period (Figure S8). As such, we found a general pattern of results that is consistent with a delay period working memory signal, which is expected to show higher confusability amongst supra-capacity set sizes. However, future work will be needed to examine changes to decoding while perfectly controlling for display-wise sensory differences. For example, it is unclear whether the poorer discriminability for high versus low set sizes during the encoding period (100-300 ms) was driven by bottom-up sensory differences or by attentional selection of items (e.g., a capacity limit in the number of selected items could contribute to the increased confusability for higher set sizes even during the early encoding period).

**Figure 6.**
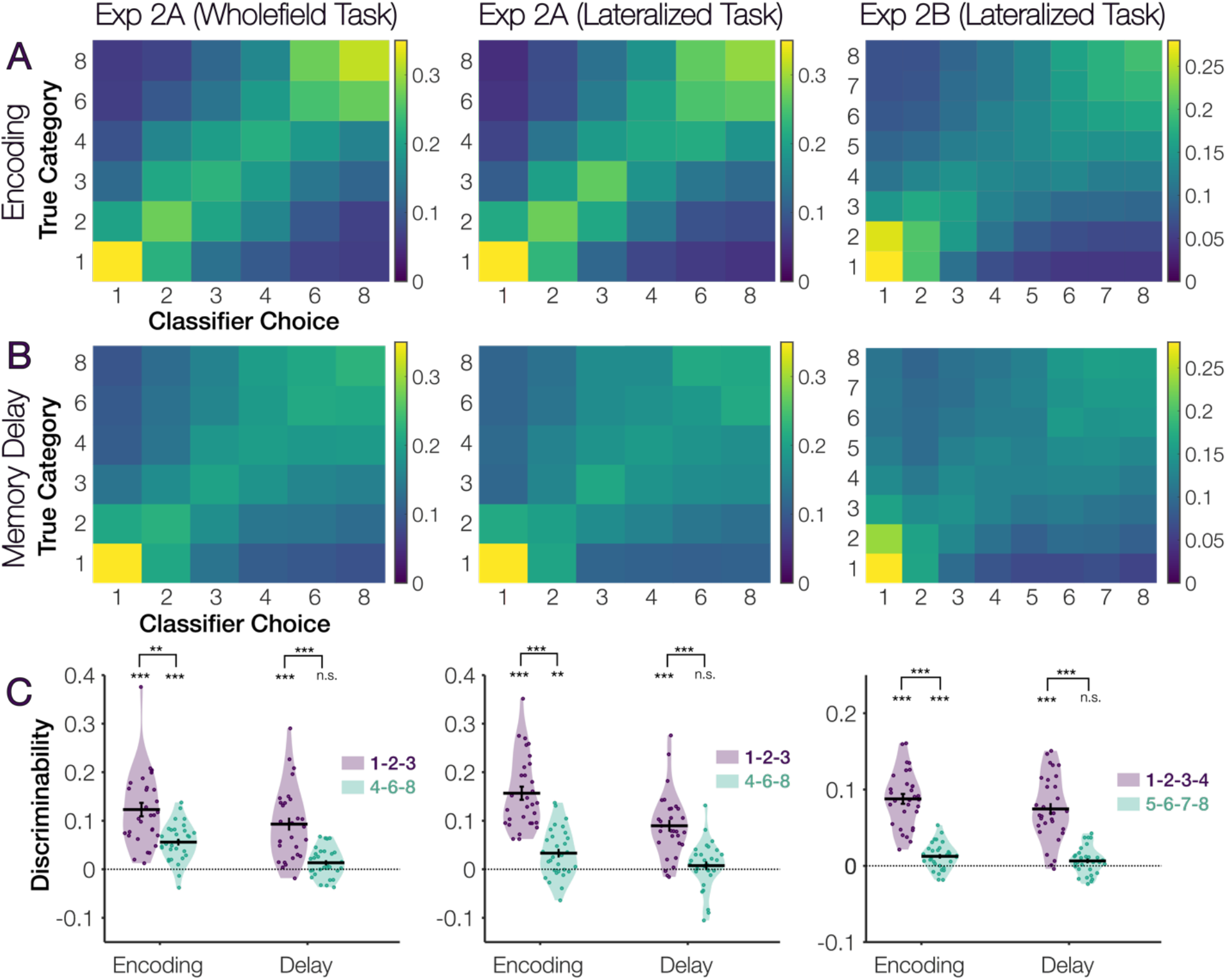
Confusion matrices and discriminability between set sizes. (A) Confusion matrix during the encoding period (100 – 300 ms). (B) Confusion matrix during the delay period (400 ms – end of delay). (C) Discriminability metric amongst lower versus higher set sizes during the encoding period and the delay period. Symbols indicate Bonferroni-corrected significance (6 comparisons), **n.s.** *p* > .10, **\*** *p* < .05, **\*\*** *p* < .01, ***\*\*\**** *p* < .001.

#### Generalization of classification across subjects and tasks

Up to this point, we have always trained and tested the classifier within a given subject. Exp 2 provided an opportunity to examine the generalizability of the classification signal across tasks, subjects, and experiments with distinct stimulus displays. In Exp 2A, the same subjects performed two different working memory task variants (one lateralized with distractors presented in the irrelevant hemifield, one whole-field display that contained no distractors). Thus, Exp 2A provided the opportunity to look at decoding within the same subject, but across distinct tasks (Figure 7A-B). Across-task decoding was overall significant during the delay period, indicating some degree of generalizability when training and testing within-versus across tasks (*p* < 1×10^−20^). However, decoding accuracy was significantly lower when training and testing across tasks versus within a task (*p* < 1×10^−9^). We quantified the difference in classification accuracy between within- and across-task decoding across 3 task epochs: encoding (100-300 ms), early delay (400 – 900 ms) and late delay (900 ms to end of delay). We found a main effect of Training (better performance for within versus across tasks), *F*(1,29) = 79.25, *p* < 1×10^−9^, η^2^_p_ = .73, and of Epoch (better performance earlier in the trial), *F*(2,58) = 31.49, *p* < 1×10^−9^, η^2^_p_ = .52. We also found an interaction of Training and Epoch, *F*(1.50,43.44) = 34.81, *p* < 1×10^− 7^, η^2^_p_ = .55, indicating that the late delay period signal was more robust to cross-generalization across task variants than the early sensory signal (i.e., during the late delay period there was no difference in classifier accuracy for within- vs. across-task training). In Figure 7C-D, we examined the ability of the classification signal to generalize across subjects within Experiment 2A, showing that there is a generalizable multivariate signature of working memory load in humans. We trained the classifier on 2/3 of data from *n-1* subjects, and tested the classifier on 1/3 of data from one held-out subject. For both the within- and across-subjects analyses, we performed the analysis separately for training within a task versus across tasks (as above). Across-subject decoding was overall significant during the delay period p < 1×10^−24^). In Figure 7C-D, we have collapsed across this task dimension but it is depicted in Figure S7. Similar to generalizing across tasks, we found that found a main effect of Training (better performance for within versus across subjects), *F*(1,29) = 45.33, *p* < 1×10^−6^, η^2^_p_ = .61, and of Epoch (better performance earlier in the trial), *F*(2,58) = 27.19, *p* < 1×10^−8^, η^2^_p_ = .48. We again found an interaction of Training and Epoch, *F*(2,58) = 44.94, *p* < 1×10^−11^, η^2^_p_ = .61, indicating that the late delay period signal was more robust to cross-generalization across subjects than the early sensory signal.

**Figure 7.**
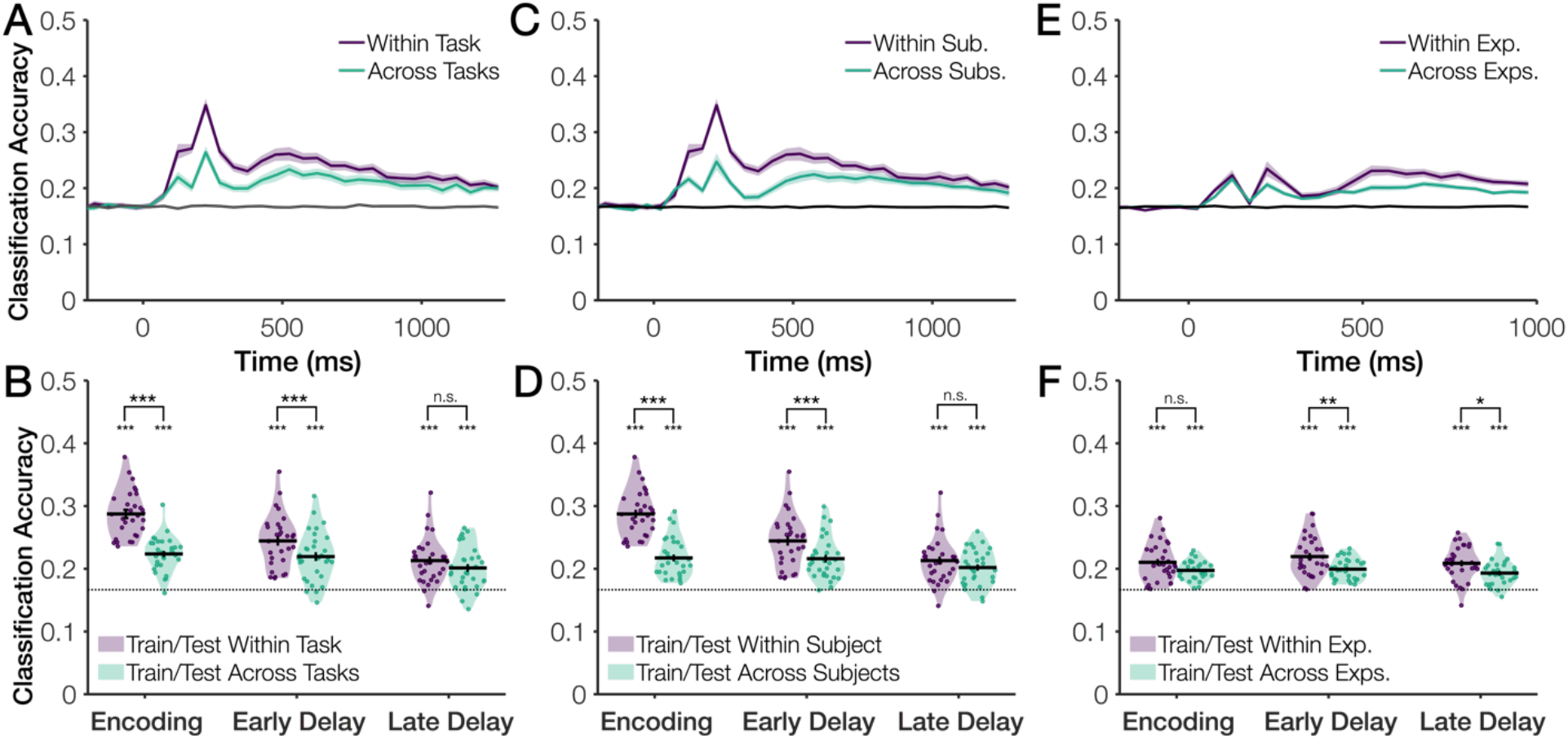
Classifier success at generalizing across task variants and across subjects. (A-B) Time-course and violin plot summaries for training/testing within a task versus across tasks (within-subjects, e.g., train on lateralized task, test on wholefield task). Shaded errors bars represent ±1 SEM. (C-D) Time-course and violin plot summaries for training/testing within subjects versus across subjects (averaged within/across tasks) (E-F) Time-course and violin plot summaries for training/testing within an experiment versus across experiments 2A and 2B (across subjects; average of within/across tasks).

Finally, in Figure 7E-F, we examined the ability of the classification signal to generalize across subjects from different experimental samples. These experiments were the same in some important ways (e.g., same electrode montage, sampling rate, and similar tasks), but differed in many minor ways (e.g., exact size of stimuli). We trained the classifier on 2/3 of data from n-1 subjects in one experiment (e.g., train Exp 2A) and tested the classifier on 1/3 of data from 1 subject in the other experiment (e.g., test Exp 2B). For both the within and across experiment analyses, we again performed the analysis separately for training within versus across tasks. Across-experiment decoding accuracy was overall significant during the delay period (*p* < 1×10^−29^). In Figure 7E-F, we have collapsed across the task dimension but it is depicted in Figure S9. We found that, relative to training across subjects but within a specific experiment, training across subjects and across experiments was slightly worse (Figure 7E-F), as indicated by a main effect of Training, *F*(1,29) = 31.33, *p* < 1×10^−5^, η^2^_p_ = .52. Here, we found no main effect of Epoch, *F*(2,58) = 1.91, *p* = .16, η^2^_p_ = .06, and no interaction of Training and Epoch, *F*(2,58) = .60, *p* = .55, η^2^_p_ = .02, indicating that the performance decrement for training across versus within experiments was consistent throughout the trial.

### Experiment 3

#### Delay-period decoding is specific to working memory task demands

In all experiments discussed so far, the amount of sensory stimulation was confounded with set size (i.e., there was more sensory stimulation on higher set size trials). Although our analyses suggest that sustained delay period decoding likely was not driven by this transient sensory confound (e.g., decoding of set sizes generalized amongst time points within the delay period, but decoding during the stimulus period did not generalize to the delay period), we wanted to test whether delay-period decoding is modulated by working memory task demands while holding visual stimulation constant. To do so, we examined data from Hakim et al. (2019). In this experiment, participants performed two different cognitive tasks using visually identical stimuli. In one condition (“Attention”) participants performed a spatial attention task. Prior work has shown that this condition did not recruit neural signatures of working memory maintenance (i.e., contralateral delay activity was absent). In the other condition (“Working memory”), participants performed a typical working memory task, and robust signatures of working memory maintenance were observed. This experiment thus provides a critical test of whether delay period decoding of set size respects the relative recruitment of working memory task demands while holding the sensory confound constant (i.e., the difference in sensory stimulation between set sizes 2 and 4 is identical for the working memory and attention task conditions).

Consistent with a signature of working memory maintenance, we observed sustained decoding of set size throughout the delay period when participants performed the working memory task, but not when they performed the spatial attention task (Fig 8A). To quantify this effect, we again divided data into the same 3 task epochs (encoding, early delay, and late delay). A repeated measures ANOVA with within subjects factors Epoch and Task revealed a main effect of Task, *F*(1.80,129.68) = 47.79, *p* < 1×10^−14^, η^2^_p_ = .40, a main effect of Epoch, *F*(1,72) = 25.90, *p* < 1×10^−5^, η^2^_p_ = .27, and an interaction of Task and Epoch, *F*(2,144) = 3.46, *p* = .034, η^2^_p_ = .05, Fig 8B. This demonstrates that early in the trial, when the sensory confound likely contributed to decoding, we observed no difference in decoding strength between the two task conditions. However, during the delay period, decoding was significantly weaker in the attention condition and completely disappeared by the late delay period (Fig 8B). This pattern of results is consistent with sustained delay period decoding as being driven by working memory task demands. To show that the null effect in the attention condition was not driven by a relatively weaker training set, we performed classification while training and testing across tasks. If the lack of decoding in the attention condition was just due to the attention condition serving poorly as a training set, then training on the working memory task should rescue delay period decoding for the attention task. However, we found no evidence of sustained delay period decoding when training across tasks. Although we were initially able to discriminate between set sizes, this early classification dissipated by around 700 ms (Fig 8C). We again performed an ANOVA with factors Epoch and Task (train Attention test WM versus train WM test Attention). We found a main effect of Epoch, *F*(1.74,125.41) = 46.55, *p* < 1×10^−13^, η^2^_p_ = .39, an effect of Task, *F*(1,72) = 6.38, *p* = .014, η^2^_p_ = .08 (slightly better overall decoding when training on the attention task, counter to the hypothetical explanation of poor attention decoding), and no interaction of Task and Epoch, *F*(2,144) = 1.19, *p* = .31, η^2^_p_ = .02 (Fig 8D). Thus, this analysis suggests that multivariate load detection is determined by storage in working memory rather than by the physical characteristics of the display, or the deployment of spatial attention alone.

**Figure 8.**
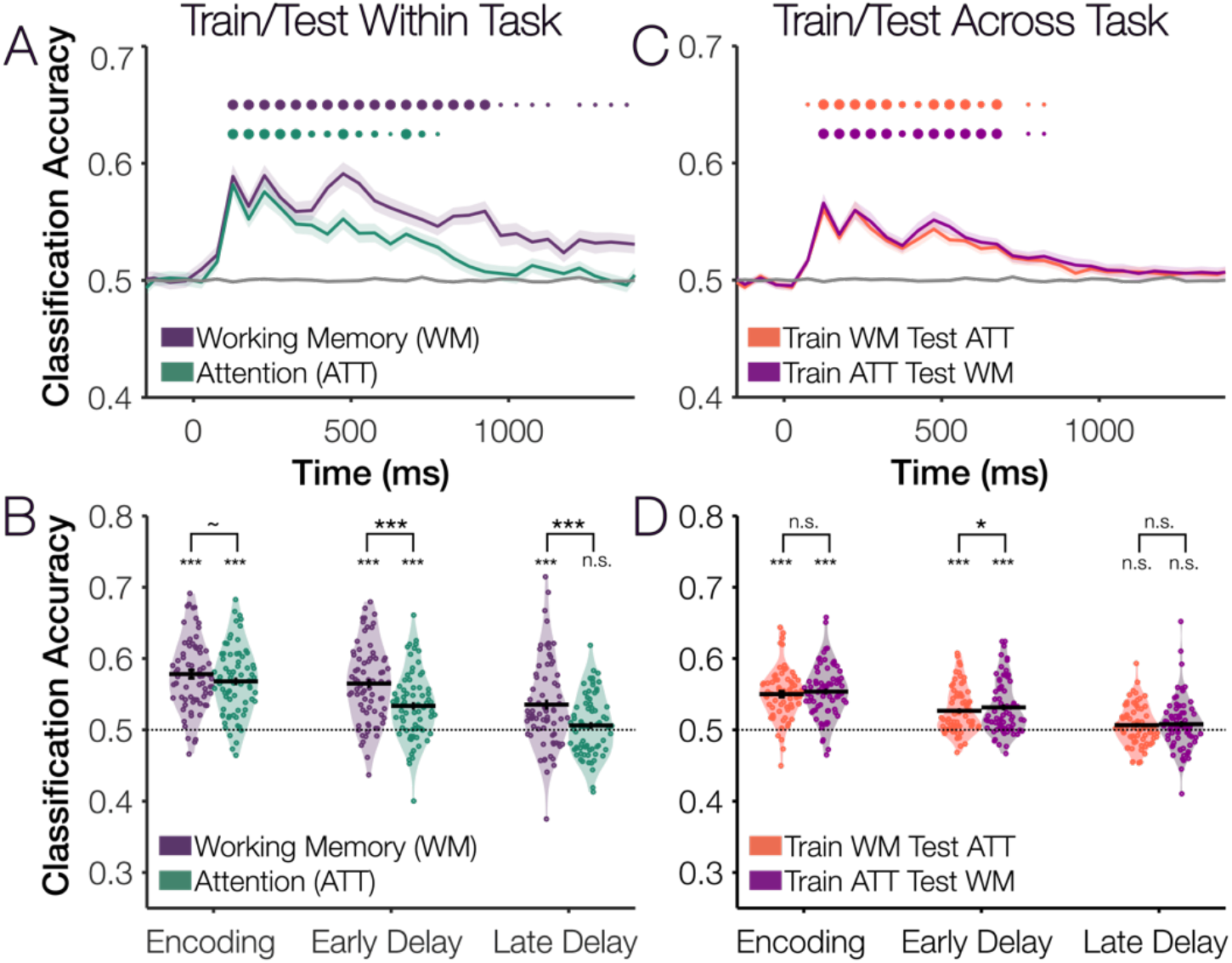
Decoding within and across tasks in Experiment 3. (A) Classification time-course for the working memory task and the lateralized attention task. Dots indicate one-tailed *t*-tests with Bonferroni-corrected significance (23 timepoints x 2 conditions = 46 comparisons), small dots, *p* < .05, medium *p* < .01, large, *p* < .001. (B) Average classification within tasks during encoding, early delay, and late delay. Stars indicate Bonferroni-corrected significance (9 comparisons), *n.s.* p > .10, ∼p <.10, *p <.05, **p <.01, ***p < .001. (C) Classification time-course training and testing across tasks (e.g., train on attention task, test on working memory task). (D) Average classification across tasks during encoding, early delay, and late delay.

### Cross-Experiment Analyses

#### Individual differences in decoding predict overall behavioral performance

Finally, we combined data from all experiments (unique subjects only) to test whether classification performance relates to behavioral performance. We reasoned that participants with higher working memory capacity would have more states to discriminate between (e.g., we would expect someone with a capacity estimate of 1.0 items to show similar neural signatures on all trials, whereas someone with a higher capacity estimate would have greater variability in delay period signatures across set size conditions). As such, we predicted that those with higher working memory capacity would likewise have higher delay period classification accuracy. Within each experiment, we z-scored behavior (average capacity, or “K”) and average delay period classification accuracy. This step was necessary to normalize differences in chance level (e.g., 50% versus 16%) and behavioral performance across experiments. Also note, we used the 10-trial ‘mini-block’ data for all four experiments. We excluded participants with capacity values lower than 2 SD’s below the group mean (i.e., who were not performing the task as instructed). We then performed a correlation using all unique subjects from all experiments. We found that classification accuracy and capacity were correlated, *r* = .26, *p* < 1×10^−4^, 95% CI = [.14, .36]. Correlations of raw classification accuracy and behavior values are shown in Figure 9B. Note, the overall correlation values did not noticeably change if we included subjects with poor behavioral performance or if we included duplicate subjects (Figure S10). Likewise, the correlation between classification accuracy and was similar in size to previously observed correlations between CDA amplitude and behavior (Figure S11). Interestingly, however, classification accuracy and CDA amplitude predicted unique behavioral variance (Analysis S1).

**Figure 9.**
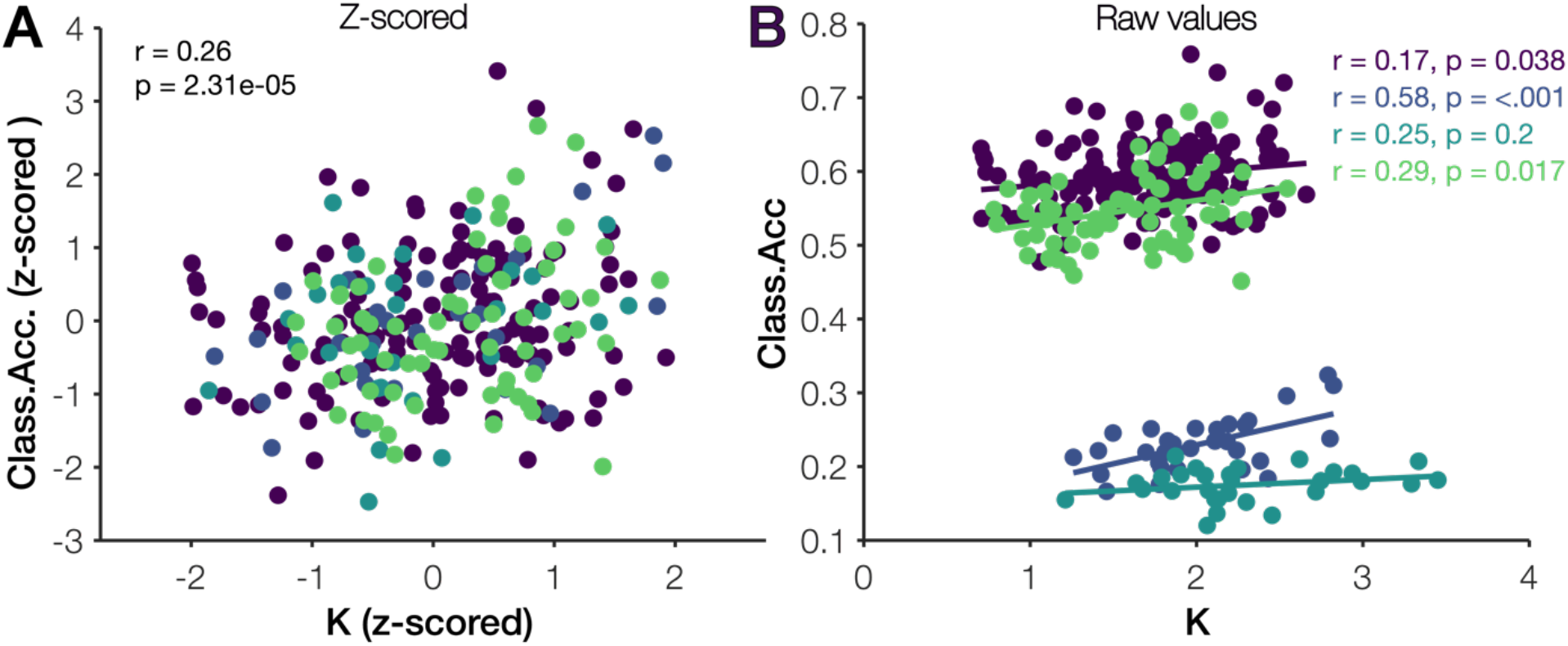
Individual differences in working memory performance and decoding accuracy. (A) Correlation between capacity and classification accuracy (z-scored) for all unique subjects in all experiments. (B) Correlations between capacity and classification accuracy (raw values) for individual experiments.

## Discussion

Here, we used a large dataset (4 published experiments, *n* = 286, >250,000 trials) to develop *multivariate load detection (“mvLoad”)*, a novel approach for tracking human working memory load using the raw EEG signal. Multivariate load detection offers several key advances over existing univariate working memory signals (e.g., contralateral delay activity, “CDA”). Most importantly, multivariate load detection is generalizable across stimulus/task differences, observers, and experiments, suggesting that it taps into a common human electrophysiological signature of working memory load. In the contexts analyzed here, it was possible to train the classifier on one set of observers, and then examine the deployment of working memory resources in a new task and set of observers. As such, multivariate load detection will allow us to study working memory in new, more flexible task contexts (e.g., without relying on lateralized displays which incur a dual task of filtering out one hemifield) and applied settings (e.g., brain-computer interfaces).

The benefits of multivariate load detection mirror similar advances made in multivariate detection of the locus of spatial attention (Rihs et al., 2007; Foster et al., 2017), attentional selection (Fahrenfort et al., 2017; Munneke et al., 2019), and an item’s visual features (Wolff et al., 2015; Bae and Luck, 2018, 2019a). Lateralized univariate EEG signals (e.g., lateralized alpha power, N2PC, CDA), have been fundamental for developing an understanding of human attention and working memory. By presenting identical visual stimuli in both hemifields, these lateralized signals exploit the contralateral organization of the human visual system. Conversely, however, to take advantage of these lateralized signals we *must* use specialized lateralized displays.

Lateralized displays offer some advantages, such as eliminating physical confounds and allowing for clever designs which place stimuli that are “invisible” to the analysis on the vertical mid-line (Hillyard et al., 1973; Hillyard and Anllo-Vento, 1998; Vogel and Machizawa, 2004; Hickey et al., 2009; Feldmann-Wüstefeld and Vogel, 2019). However, lateralized CDA designs also introduce potential disadvantages. First, when presenting to-be-remembered items in both hemifields, there is some ambiguity as to whether differences in the CDA and behavior are confounded by the joint need to suppress irrelevant visual information. With increasing memory set size, there is both an increased need to remember more information and an increased amount of irrelevant visual information to suppress. Second, to measure lateralized components such as the CDA, we must construct a difference score (contralateral – ipsilateral). The statistical reliability of difference scores is poor when the two underlying measures are highly correlated (Rodebaugh et al., 2016). Due to the poor spatial resolution of EEG, trial-by-trial voltage scores for contralateral and ipsilateral electrodes are highly correlated, thus the reliability of single-trial difference scores is lower than from single electrodes. Although we still obtain reliable estimates of CDA amplitude when averaging across many hundreds of trials, this traditional univariate approach potentially throws away valuable single-trial information that could be exploited to better predict working memory load.

In the current work, we used several datasets to demonstrate the utility of multivariate load detection in many contexts. We consistently found robust, sustained decoding of working memory load throughout the memory delay period, and this decoding predicted individual differences in working memory behavior. Multivariate load detection was sensitive to fine-grained variations in memory load as well as to working-memory specific (as opposed to general attentional) task demands. Further, we showed that multivariate load detection generalized across stimulus differences (e.g., remembering colors versus shapes; lateralized versus whole-field presentation of the items) and generalized across observers (e.g., we can train the decoder on a large group of subjects, then predict memory load in a new subject whose data the classifier has never seen). The high generalizability, in particular, will be critical for future work; using the approach outlined here, we think it is possible to build a generalizable, pre-trained classifier which will be able to predict visual working memory load using relatively small amounts of data from new tasks and observers.

Future work will need to address some potential limitations of the current work. First, because we used previously published datasets, we were limited in our ability to perfectly control for potential confounds such as visual stimulation (i.e., transient luminance changes also increased with memory load). Future work using manipulations such as selective encoding (i.e., holding visual stimulation constant but varying which items are encoded) or retro-cues (Griffin and Nobre, 2003; Lepsien and Nobre, 2007; Harrison and Tong, 2009; Christophel et al., 2012, 2018; Sprague et al., 2016), will be critical for disentangling encoding-related decoding from bottom-up, visually-driven decoding of load during the early part of the trial. Second, a key contribution of this work is its demonstration of the feasibility of building a generalizable, pre-trained classifier for detecting working memory load in new tasks and observers. Here, we demonstrate that this type of generalizability is feasible across some stimulus differences and across observers within a site (i.e., controlling for high-level differences in experimental procedures such as amplifier, referencing, and montage). Future work will be needed to further investigate the cross-site generalizability of multivariate load detection (e.g., variations in EEG systems, reference, experimental procedures, and subject pools) to build a generalizable, pre-trained working memory load detector (Dansereau et al., 2017; Scheinost et al., 2019).

Finally, we anticipate that this method will also be useful outside of EEG. In the fMRI literature, for example, prior work has found univariate, load-dependent changes in parietal and prefrontal cortex (Braver et al., 1997; Cohen et al., 1997; Todd and Marois, 2004, 2005; Xu and Chun, 2006). In early visual cortex, in contrast, there are no load-dependent changes to the univariate signal with load. Despite this, the identity of a single item can be robustly decoded from visual cortex (Kamitani and Tong, 2005; Harrison and Tong, 2009; Serences et al., 2009). Further, the fidelity of item-specific decoding in visual cortex is degraded with load (Emrich et al., 2013; Sprague et al., 2014). However, no extant work has looked at multivariate neural signatures of load in visual cortex for supra-capacity set sizes, in part because of limitations of the spatial resolution of the method (with more than 2-3 items, the voxel population receptive fields would start to overlap substantially). Multivariate load detection, as performed here, could be used to probe whether working memory load, per se, can be decoded in visual cortex. Further, this method could be used to test whether long-observed univariate signatures in parietal and frontal cortex are purely univariate in nature, or if more information about memory load can be gleaned by applying multivariate methods.

We argue that multivariate load detection (“mvLoad”) is a generalizable electrophysiological marker of human working memory load, and that this approach will allow for the unprecedented combination of disparate datasets to build a powerful, generalizable model of human working memory load. Because multivariate load detection is generalizable across tasks and observers, we anticipate that this method will be useful in many basic and applied research settings (i.e., unobtrusively monitoring the contribution of working memory during other cognitive contexts). All data shown here are available on the Open Science Framework (upon publication). We encourage other labs using distinct populations (e.g., developmental; clinical), research sites (e.g., outside of the U.S.) and task variants to use our published data to test the extent of the generalizability of this new method.

## Data availability

Data for all experiments are freely available the Open Science Framework at https://osf.io/6jkqu/

## Acknowledgements

We thank our co-authors on the published datasets used to conduct this work: Keisuke Fukuda, Eren Gunseli, Nicole Hakim, Irida Mance, Nash Unsworth & Geoff Woodman.

## Footnotes

* Greenhouse-Geisser correction applied when the assumption of sphericity is violated

