## Supporting Information for "Multivariate analysis reveals a generalizable human electrophysiological signature of working memory load"

### Supplemental Methods

Additional details about the EEG acquisition parameters and pre-processing of the EEG data in each of the published papers. Note, some text about the methods is quoted from the original published papers.

**Experiments 1, 2A and 2B: Unsworth, Fukuda, Awh & Vogel (2015); Fukuda, Mance & Vogel (2015); Fukuda, Woodman, Vogel (2015).**

Participants were seated in an electrically shielded chamber with heads resting on a padded chin rest ~100 cm the monitor (17-in CRT, refresh rate = 60 Hz). EEG data were collected from 20 passive tin electrodes (SA Instrumentation Co., San Diego, CA) mounted in an elastic cap (ElectroCap International, Eaton, OH). Electrode positions included International 10/20 sites F3, Fz, F4, T3, C3, Cz, C4, T4, P3, Pz, P4, T5, T6, O1, and O2 and five non-standard sites: OL midway between T5 and O1, OR midway between T6 and O2, PO3 midway between P3 and OL, PO4 midway between P4 and OR, POz midway between PO3 and PO4. Data were recorded with a left-mastoid reference and re-referenced offline to the algebraic average of the left and right mastoid. Horizontal electrooculogram (EOG) and vertical EOG were collected from 3 additional passive electrodes affixed to the face with stickers. The EEG and EOG were amplified with an SA Instrumentation amplifier with a bandpass of 0.01– 80 Hz and were digitized at 250 Hz using LabVIEW 6.1 running on a PC.

Trials were epoched before and after the memory array onset (Experiment 1: -200 to 996 ms; Experiment 2A: -500 to 1300 ms; Experiment 2B: -500 to 1048 ms). and baselined to the pre-trial period (Experiment 1: -200 to 0 ms; Experiment 2A and B: -500 to 0 ms). In Experiment 1, trials were visually inspected for artifacts, and discarded trials contaminated by blocking, blinks, detectable eye movements, excessive muscle noise, or skin potentials were discarded. Here, we used the same artifact index (i.e., a vector of 0's and 1's marking whether an artifact is present or absent for each trial) that was originally used in Unsworth et al. 2015. In Experiments 2A and 2B, we re-identified artifacts using previously published procedures, and manually inspected trial waveforms that the procedures worked as expected. For HEOG rejection, we used a split-half sliding-window approach (window size = 100 ms, step size = 10 ms, threshold = 12 µV). We slid a 100-ms time window in steps of 10 ms from the beginning to the end of the trial. If the change in voltage from the first half to the second half of the window was greater than 12 µV, it was marked as an eye movement and rejected. For blinks, we used a sliding-window step function on data from the vertical EOG (window size = 100 ms, step size = 10 ms, threshold = 15 µV). Note, these HEOG and VEOG thresholds are more strict than those used in Experiment 3 (below) because of (1) slightly different scaling of data from the passive amplifier (as data from this amplifier were manually scaled to reference pulses) and (2) lack of eye tracking data to complement the EOG. To check for drift, we compared the absolute change in voltage from the first quarter of the trial to the last quarter of the trial. If the change in voltage exceeded 100 $\mu$V, the trial was rejected for drift. In addition to slow drift, we checked for sudden step-like changes in voltage with a sliding window (window size = 250 ms, step size = 20 ms, threshold = 100 $\mu$V). We excluded trials for muscle artifacts if any electrode had peak-to-peak amplitude greater than 200 $\mu$V within a 15-ms time window. We excluded trials for blocking if any electrode had at least 30 time points in any given 200-ms time window that were within 1$\mu$V of each other.

**Experiment 3: Hakim, Adam, Gunseli, Awh & Vogel (2019)**

**Acquisition.** Participants were seated in an electrically shielded chamber with heads resting on a padded chin rest ~74 cm from the monitor (24-in LCD, refresh rate = 120 Hz). Data were collected from 30 active Ag/AgCl active electrodes (actiCHamp, Brain Products, Munich Germany) mounted in an elastic cap positioned according to the international 10-20 system (Fp1, Fp2, F7, F8, F3, F4, Fz, FC5, FC6, FC1, FC2, C3, C4, Cz, CP5, CP6, CP1, CP2, P7, P8, P3, P4, Pz, PO7, PO8, PO3, PO4, O1, O2, Oz). Two additional active electrodes were affixed with stickers to the left and right mastoids, and a ground electrode was placed at position Fpz. Data were referenced online to the right mastoid and re-referenced offline to the algebraic average of the left and right mastoids. Incoming data were filtered (low cutoff = .01 Hz, high cutoff = 80 Hz; slope from low to high cutoff = 12 dB/octave) and recorded with a 500-Hz sampling rate. Impedance values were kept below 10 kΩ.

Eye movements and blinks were monitored using electrooculogram (EOG) activity and eye tracking. We collected EOG data with five passive Ag/AgCl electrodes (two vertical EOG electrodes placed above and below the right eye, two horizontal EOG electrodes placed ~1 cm from the outer canthi, and one ground electrode placed on the left cheek). We collected eye- tracking data using a desk-mounted EyeLink 1000 Plus eye-tracking camera (SR Research, Ontario, Canada) sampling at 1,000 Hz. Usable eye-tracking data were acquired for 90 participants.

**Artifact rejection and pre-processing.** Data were segmented into trials (900 ms before memory array onset to 1,450 ms after memory array onset) and trials were baselined to the period -700 ms to -400 ms before the memory array onset (this baseline was chosen to avoid the cue onset which happened at around 300 ms before memory array onset). Trials containing ocular artifacts, movement artifacts, blocking, or drift were excluded from analyses. Artifacts were first detected via automatic criteria, and then trials were manually inspected to ensure criteria were working as expected. To detect eye movements, we used a sliding-window step function on data from the horizontal EOG (HEOG) and the eye-tracking gaze coordinates. For HEOG rejection, we used a split-half sliding-window approach (window size = 100 ms, step size = 10 ms, threshold = 20 µV). We used the HEOG rejection only if the eye-tracking data were bad for that trial epoch. We slid a 100-ms time window in steps of 10 ms from the beginning to the end of the trial. If the change in voltage from the first half to the second half of the window was greater than 20 µV, it was marked as an eye movement and rejected. For eye-tracking rejection, we applied a sliding-window analysis to the x-gaze coordinates and y-gaze coordinates (window size = 100 ms, step size = 10 ms, threshold = 0.5° of visual angle). For blinks, we used a sliding-window step function on data from the vertical EOG (window size = 80 ms, step size = 10 ms, threshold = 30 µV). We checked the eye- tracking data for trial segments with missing data points (no position data are recorded when the eye is closed). To check for drift, we compared the absolute change in voltage from the first quarter of the trial to the last quarter of the trial. If the change in voltage exceeded 100 $\mu$V, the trial was rejected for drift. In addition to slow drift, we checked for sudden steplike changes in voltage with a sliding window (window size = 100 ms, step size = 10 ms, threshold = 100 $\mu$V). We excluded trials for muscle artifacts if any electrode had peak-to-peak amplitude greater than 200 $\mu$V within a 15-ms time window. We excluded trials for blocking if any electrode had at least 30 time points in any given 200-ms time window that were within 1$\mu$V of each other.

### Supplemental Results

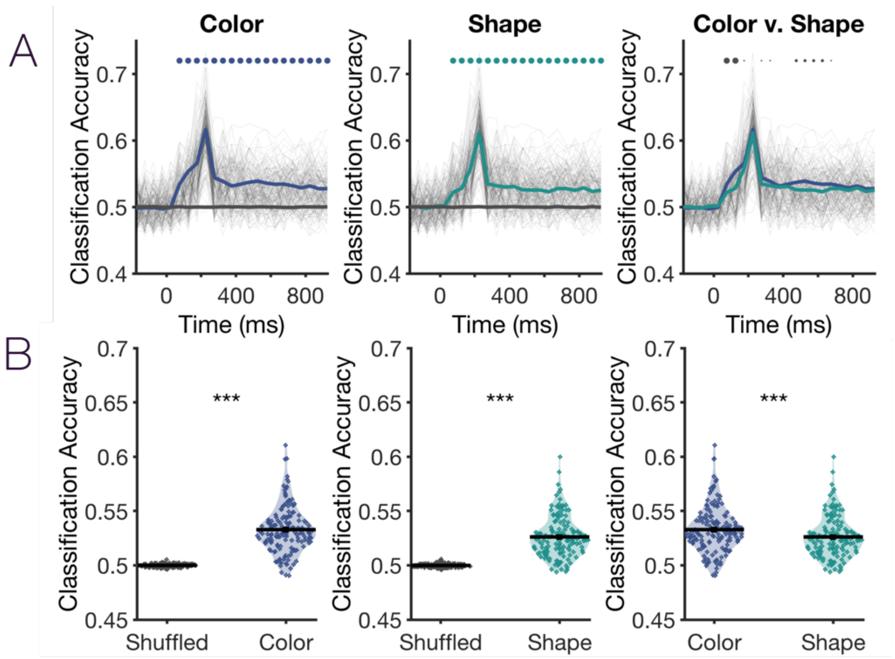


**Figure S1. Classification separated by the color and shape conditions.** Analyses in Figure 2 (Experiment 1) were performed on data combined across two conditions: remembering colors and remembering shapes. Here, we show that single-trial classification results are qualitatively similar when performed on color or shape data alone. (A) Time course of classification for color and shape conditions. Dots represent Bonferroni corrected significance for 23 time bins (small dots $p <.05$, medium $p < .01$, large $p < .001$). (B) Overall delay period classification. Both shape and color classification were robustly above chance (shuffled baseline), though it was slightly higher for the color condition compared to the shape condition.

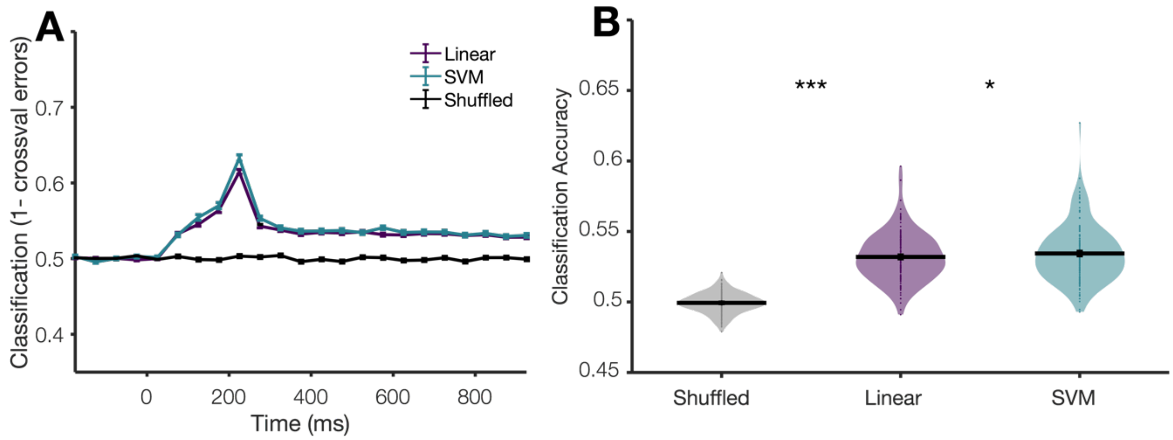


**Figure S2. Linear classifier performance versus a support vector machine.** (A) Classification performance over time with a standard linear classifier versus a support vector machine. The support vector machine was fit and cross-validated with Matlab functions *fitcsvm.m* (options: standardized predictors; Gaussian kernel function, auto kernel scale), *crossval.m* (default options), and *kfoldLoss.m* (default options). (B) Average classification performance during the delay period ($p_{uncorrected}$ = .01). The SVM yielded a modest improvement over the linear classifier (but, offered less speed and flexibility than the linear classifier we chose).

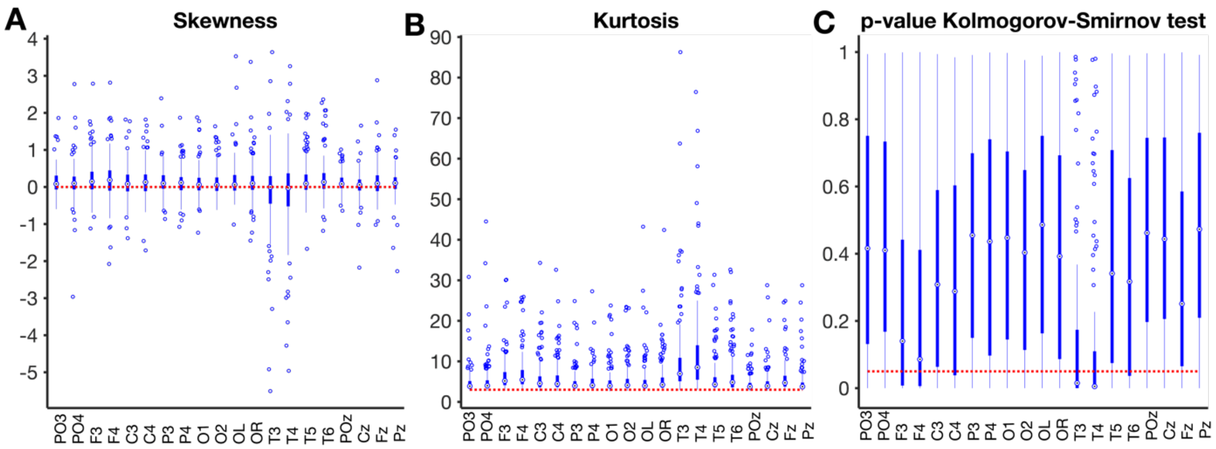


**Figure S3. Normality of trial voltage distributions at each electrode.** The relatively modest performance difference between the linear classifier and the SVM suggests that our data not violate the linear classifiers' assumption of normality. To test this, we calculated skewness, kurtosis, and ran a Kolmogorov-Smirnov test on each iteration of the analysis. These calculations were done separately for each electrode, but on one average value for the delay period (i.e., averaged across time points 400-1000 ms) for all trials (i.e., a vector of all trials from all set size conditions). For our nTrials x nElectrodes matrix, we tested whether each electrode had normally or non-normally distributed trial voltage values on each iteration of the classification analysis. We repeated this analysis for every participant and every random iteration of trial assignment to the training set. We found that the trial distributions were numerically very slightly skewed (Median skewness = 0.09) and moderately leptokurtic (Median kurtosis = 4.41). However, the trial distributions for most electrodes were not significantly different from normal as assayed by the Kolmogorov-Smirnov test (with the exception of T3 and T4, which were the most strongly leptokurtic and had modal p-values below .05). (A) Box plots of the skewness of trial voltages at each electrode. In the box plots, the middle bullseye represents the median value, the thick blue bar represents values in the $25^{th}$ to $75^{th}$ percentile, and the thin blue lines ("whiskers") represent the range of values not considered outliers. Outliers are plotted as dots outside the whiskers. A normal distribution is expected to have a skewness value of 0 (red dotted line). (B) Box plots of the kurtosis of trial voltage distributions at each electrode. A normal distribution is expected to have a kurtosis value of 3 (red dotted line). (C) Box plots of *p*-values from the Kolmogorov-Smirnov test at each electrode show that p-values are broadly distributed in the null range, with the median p-value falling below p = .05 (red dotted line) for only electrodes T3 and T4 (each box plots shows the distribution of values at each electrode for all the iterations and subjects in the analysis).

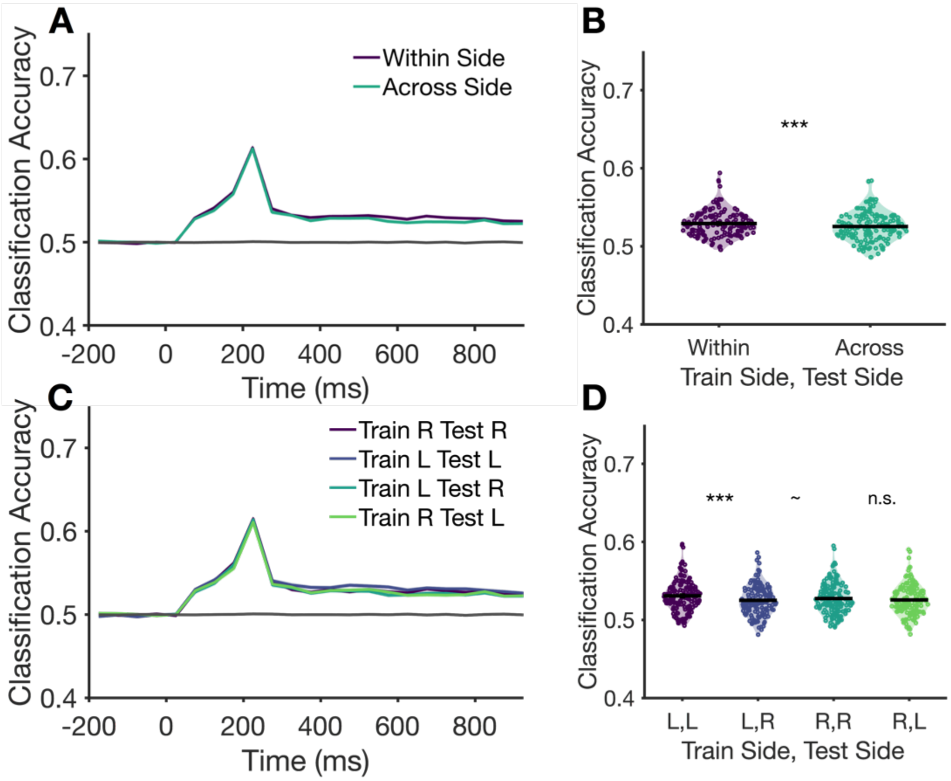


**Figure S4. Classification within versus across cued sides in Experiment 1.** (A) Classification accuracy over time as a function of training/testing within a side (e.g., train on "remember left", test on "remember left") versus across cued sides (e.g., train on "remember left", train on "remember right"). (B) Average delay period decoding for training/testing within versus across sides. (C) Classification of accuracy over time, with separate lines shown for all possible combinations of training/testing on each side. (D) Average delay period decoding for data shown in (C). The within-side decoding benefit was significant for left side, but not right side, training data. *** p<.001, ~ p<.10, n.s., p>.10.

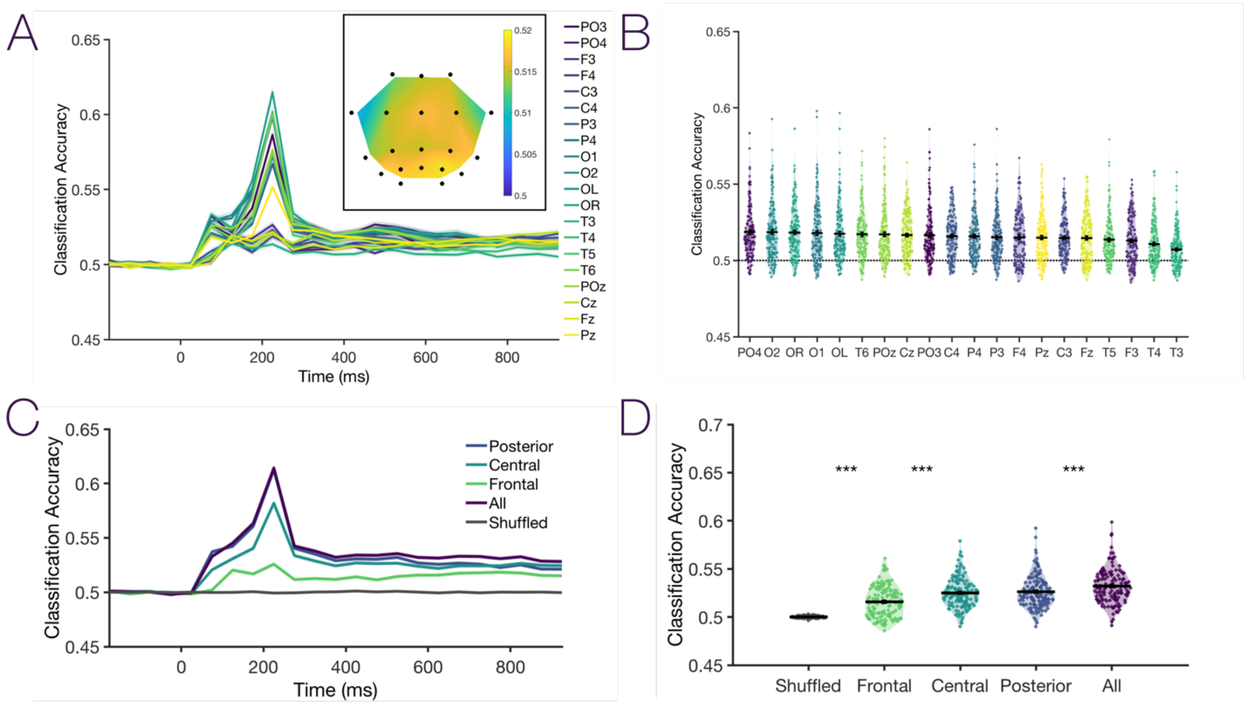


**Figure S5. Classification for single electrodes and electrode groups.** (A) We ran the single-trial classification analysis separately for each electrode (i.e., using only 1 predictor). The main figure shows classification accuracy over time for each electrode. The inset shows a topographic plot of average decoding accuracy during the delay period. (B) Average decoding accuracy during the delay period. All electrodes significantly outperformed a shuffled baseline ($p$ < .001 Bonferroni-corrected for 20 channels). Electrodes are plotted from left to right in relative descending order of classification accuracy. (C). We likewise ran the single-trial classification analysis separately for broad groups of electrodes, including posterior (O1,O2,OL,OR,PO3,PO4,T5,T6), central (C3,Cz,C4,P3,Pz,P4,T3,T4), and frontal (F3,F4,Fz). (D) Average classification accuracy for groups of trials electrodes during the delay period. All groups significantly predicted set size, though frontal electrodes were had lower classification accuracy than central or posterior electrodes. However, no single group of electrodes achieved the same classification accuracy as the full analysis (all 20 electrode predictors). Together with Figure 3, these supplementary analyses suggest that changes to voltage values across the entire scalp (i.e., changes to the degree of positivity at frontal sites and changes to the degree of negativity at the posterior cites) are likely driving overall decoding performance.

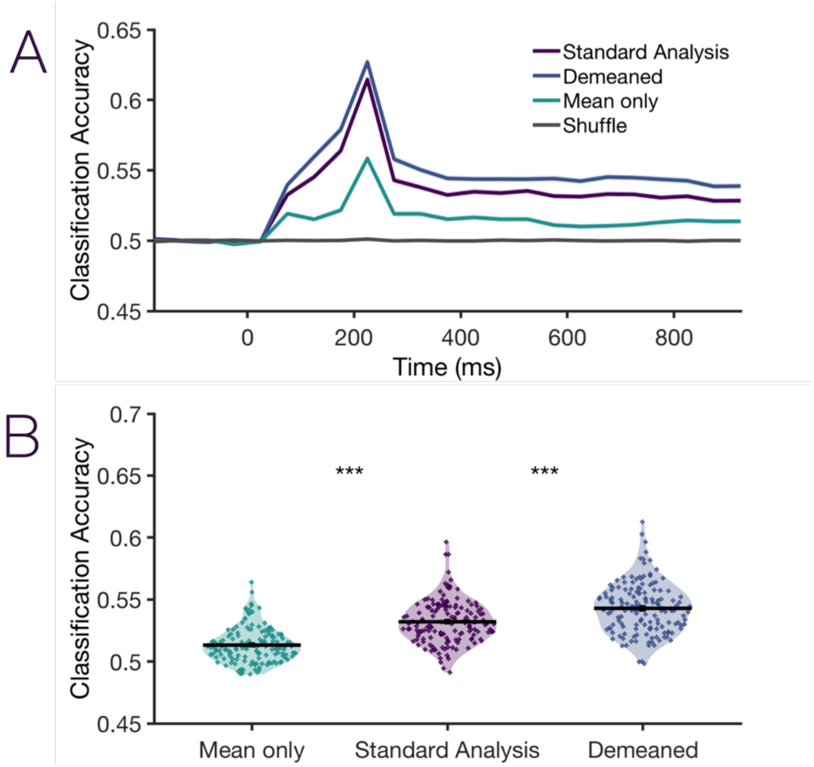


**Figure S6. Classification using the global signal or the de-meaned pattern across electrodes.** Given that even single electrodes (e.g., 1 predictor as in Figure 2) predicted working memory load, we wanted to test whether the multivariate aspect of this approach is truly beneficial or if, instead, the data could be equally well accounted for by a simple univariate signal change. To test this, we compared our standard analysis (raw voltage values from 20 electrode predictors) with demeaned values (i.e., take subtract the trial-wise mean from the 20 electrode predictors) and with the mean univariate signal alone (i.e., use the global univariate signal as our only predictor, quantified as 1 average voltage value for all electrodes). Overall, we found that the multivariate aspect of this signal (i.e., the pattern of voltage across electrodes) had more predictive power than a univariate signal alone. Although the univariate signal predicted working memory load significantly above chance ($p < .001$), it was significantly less predictive than the standard analysis ($p < .001$) which was in turn less effective than the trial-wise demeaned analysis ($p < .001$). (A) Time course of classification accuracy. (B) Violin plots of mean classification accuracy during the delay period.

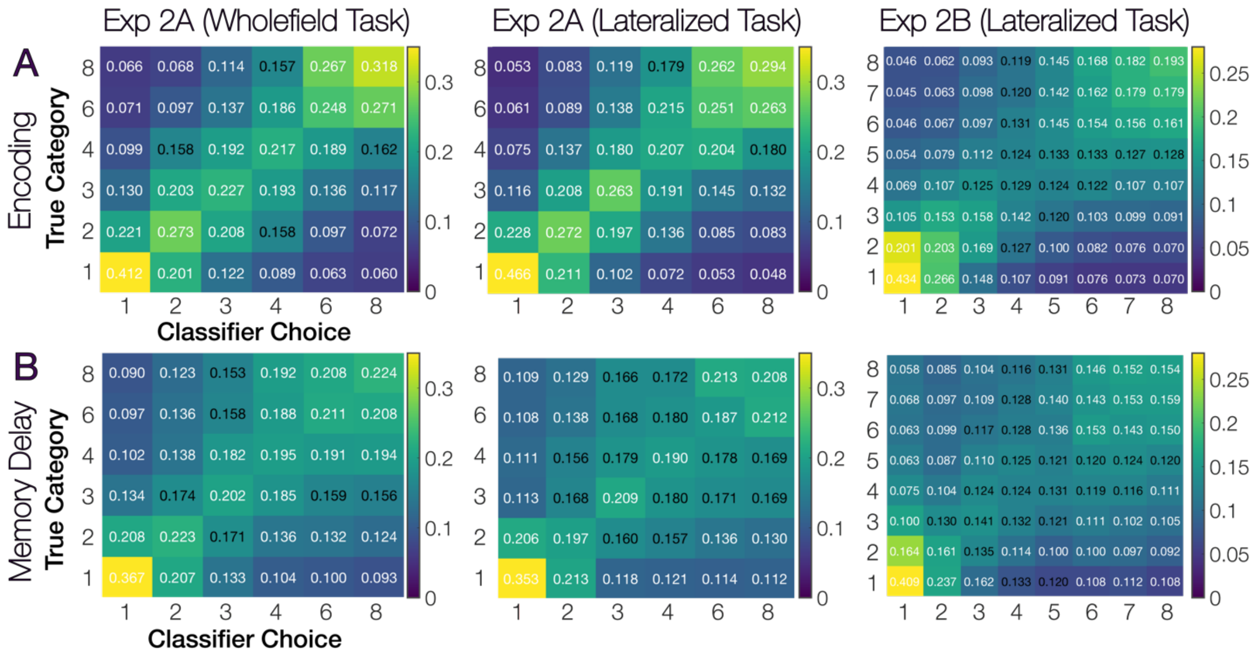


**Figure S7. Appendix of numerical values for each cell of the confusion matrix plots.** Text values in each cell show the numerical proportion values for each cell in the confusion matrix. White text indicates that the cell was significantly different from (either higher or lower than) the corresponding cell in a confusion matrix made from the permuted data ($p_{uncorrected}$). For a more direct comparison of decoding each set size relative to others, please see Figure S8 for pairwise classifiers of all possible pairs of set sizes in each experiment.

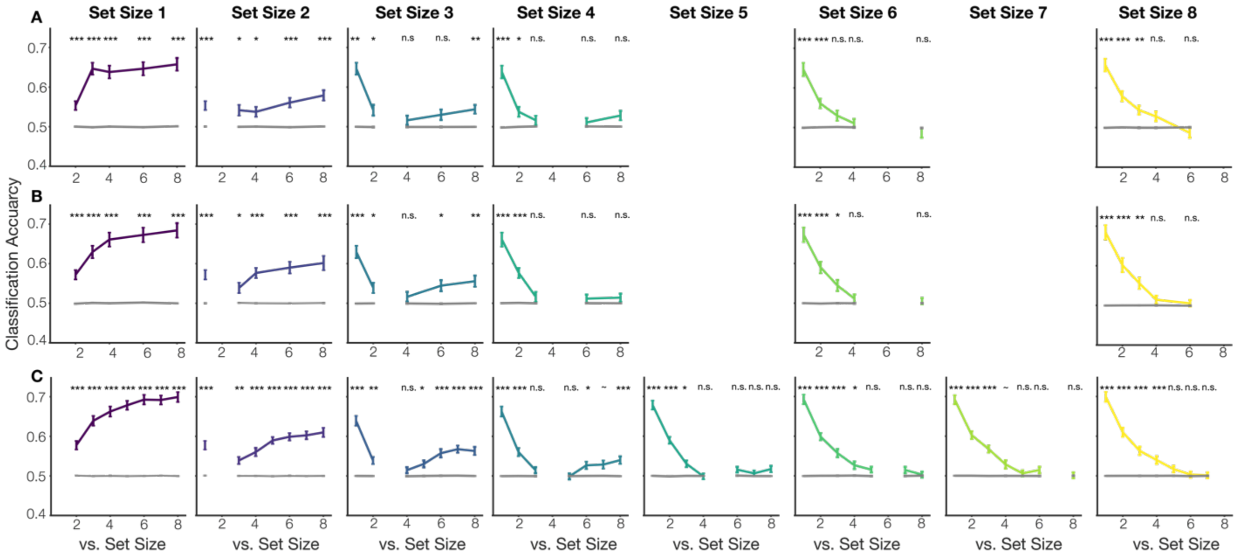


**Figure S8. Pairwise classifiers during the delay period.** (A) Pairwise classifiers for the whole-field task in Experiment 2A. Each subplot shows the classification performance when doing pairwise classification between the set size in the title of the column (e.g., 1st column = set size 2) pitted against every other possible set size (x-axis). (B) Pairwise classifiers for the lateralized task in Experiment 2A. (C) Pairwise classifiers for Experiment 2B (also a lateralized task). Set Sizes 1-3 were most discriminable from one another and from all other set sizes. Set Sizes 4-8 were highly confusable with one another. *** p<.001, **p<.01, *p<.05, ~p<.10, n.s. p >.10 (Comparisons are bonferroni-corrected for the number of comparisons within each sub-plot).

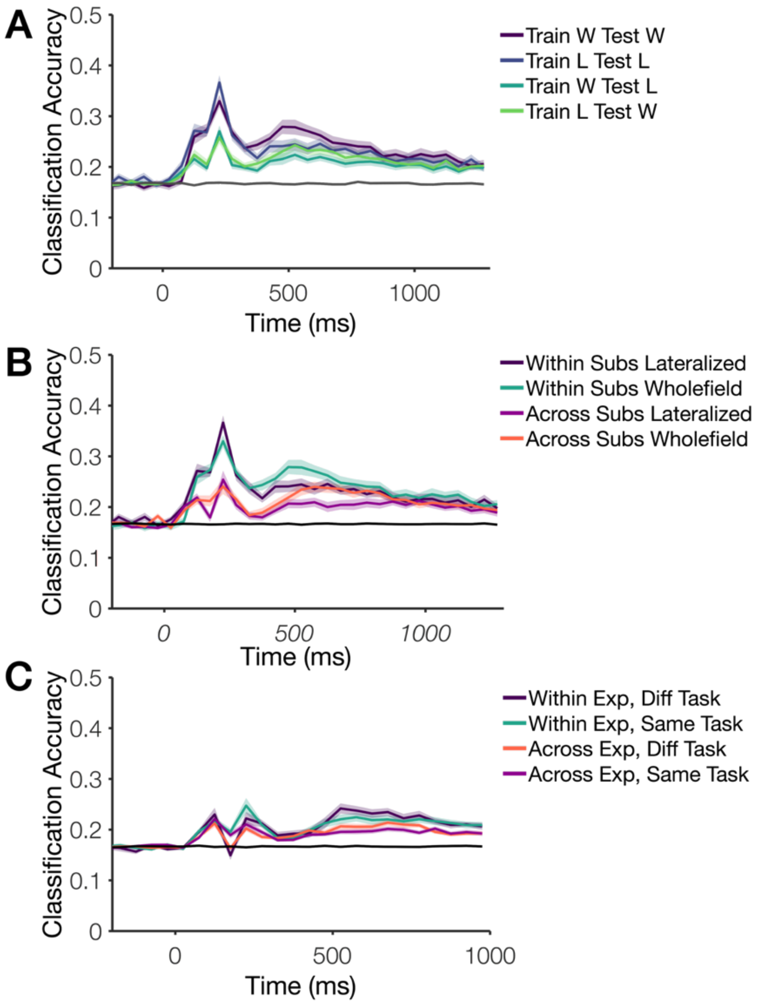


**Figure S9. All combinations for cross-task, -subject, and -experiment analyses.** (A) Training within and across tasks (within subjects) in Experiment 2A. W is short for the wholefield task, and L is short for the lateralized task. In the main text, the line “within task” is the average of “Train L Test L” and “Train W Test W” lines; the “across tasks” line is the average of the “Train L Test W” and the “Train W Test L” lines. (B) Training within and across subjects in Experiment 2A. We performed the within and across subjects analysis separately for the lateralized and wholefield tasks. In the main text, the line “Within Subs” is the average of the lines “Within Subs Lateralized” and “Within Subs Wholefield” in this figure; the line “Across Subs” is the average of the lines “Across Subs Wholefield” and “Across Subs Lateralized” in this figure. (C) Training within and across experiments (2A and 2B). We performed the within and across experiments analysis separately while also training/testing within or across tasks. In the main text, the line “Within Exp” is the average of the lines “Within Exp, Diff Task” and “Within Exp, Same Task” in this figure; the line “Across Exps.” is the average of the lines “Across Exp, Same Task” and “Across Exp, Diff Task” in this figure.

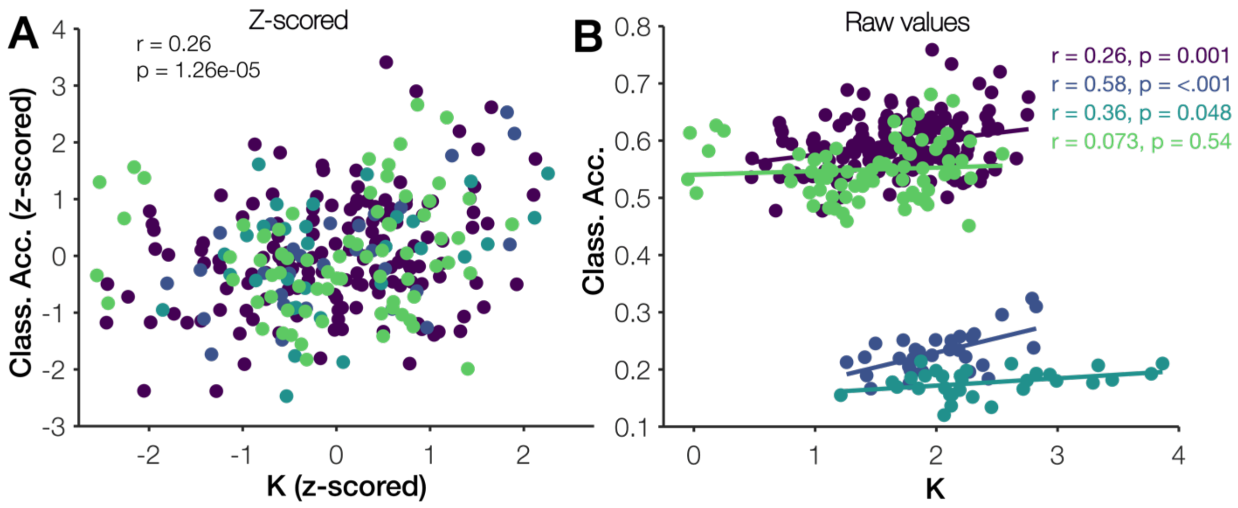


**Figure S10. Individual differences in classification accuracy and working memory behavior (K) all unique subjects (no subjects excluded for poor performance, > 2 S.D.'s below group average).** (A) Combined correlation using z-scored behavior and classification accuracy data. (B) Individual experiment correlations, including all unique subjects.

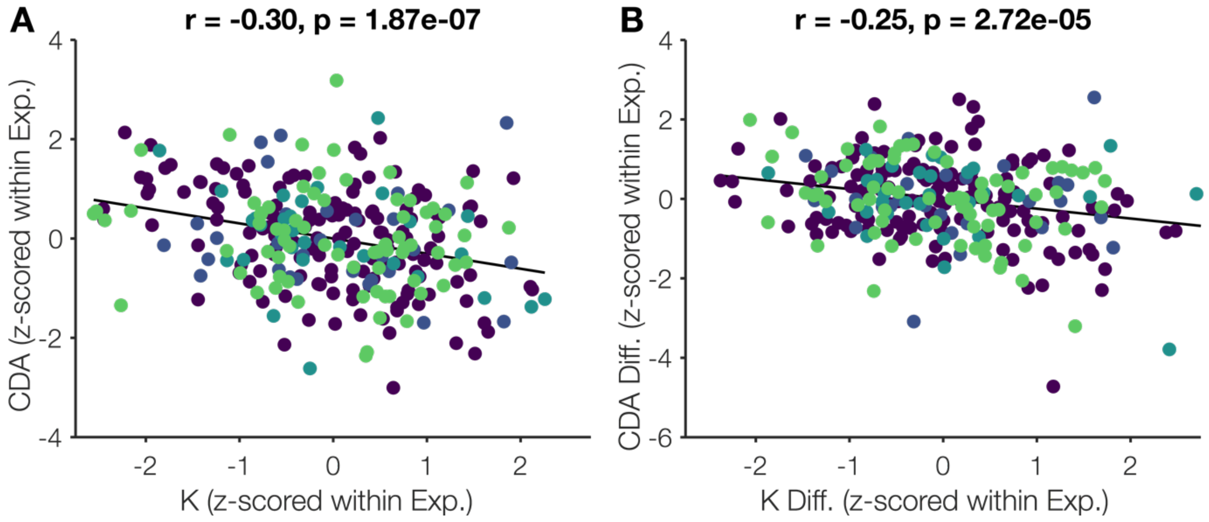


**Figure S11. Individual differences in CDA amplitude and working memory behavior (K) for all unique subjects (no subjects excluded for poor performance).** (A) Combined correlation using z-scored behavior (K) and z-scored CDA amplitude (average of all set sizes). (B) Combined correlation using a z-scored behavioral difference (average of higher set sizes minus lower set sizes) and z-scored CDA difference (average of higher set sizes minus lower set sizes). Correlation values were quite similar to those observed for the relationship between classification accuracy and behavioral performance.

**Analysis S1.** Correlation values were quite similar to those observed for the relationship between classification accuracy and behavioral performance. However, a stepwise linear regression revealed that classification accuracy, overall CDA amplitude and the CDA set size difference predicted unique variance in behavior. Using z-scored data from all participants across all experiments, we ran a step-wise linear regression. Model 1 (Average CDA amplitude) predicted 9.3% of the variance (R = .304, $p < .001$). Model 2 (Average CDA, + Classification accuracy) predicted significantly more variance (R = .358, $\Delta R^2$ $p < .001$). Model 3 likewise predicted significant additional variance (R = .389, $\Delta R^2$ $p = .007$).

**Model Summary**

| Model | R | $R^2$ | Adjusted $R^2$ | RMSE | $R^2$ Change | F Change | df1 | df2 | p |
|---|---|---|---|---|---|---|---|---|---|
| 1 | 0.000 | 0.000 | 0.000 | 0.995 | 0.000 | | 0 | 281 | |
| 2 | 0.304 | 0.093 | 0.089 | 0.949 | 0.093 | 28.573 | 1 | 280 | 1.874e-7 |
| 3 | 0.358 | 0.128 | 0.122 | 0.932 | 0.036 | 11.386 | 1 | 279 | 8.448e-4 |
| 4 | 0.389 | 0.151 | 0.142 | 0.921 | 0.023 | 7.468 | 1 | 278 | 0.007 |

**Coefficients**

| Model | | Unstandardized | Standard Error | Standardized | t | p |
|---|---|---|---|---|---|---|
| 1 | (Intercept) | -3.616e-7 | 0.059 | | -6.105e-6 | 1.000 |
| 2 | (Intercept) | 5.147e-8 | 0.057 | | 9.107e-7 | 1.000 |
| | ZCDAall | -0.304 | 0.057 | -0.304 | -5.345 | 1.874e-7 |
| 3 | (Intercept) | -2.932e-7 | 0.056 | | -5.283e-6 | 1.000 |
| | ZCDAall | -0.257 | 0.058 | -0.257 | -4.463 | 1.176e-5 |
| | ClassZ | 0.194 | 0.058 | 0.194 | 3.374 | 8.448e-4 |
| 4 | (Intercept) | -4.099e-7 | 0.055 | | -7.471e-6 | 1.000 |
| | ZCDAall | -0.220 | 0.059 | -0.220 | -3.748 | 2.164e-4 |
| | ClassZ | 0.193 | 0.057 | 0.193 | 3.380 | 8.279e-4 |
| | ZCDAdiff | -0.156 | 0.057 | -0.156 | -2.733 | 0.007 |
